# Laminarin-triggered defence responses are geographically dependent for natural populations of *Solanum chilense*

**DOI:** 10.1101/2021.06.25.449942

**Authors:** Parvinderdeep S. Kahlon, Andrea Förner, Michael Muser, Mhaned Oubounyt, Michael Gigl, Richard Hammerl, Jan Baumbach, Ralph Hückelhoven, Corinna Dawid, Remco Stam

**Affiliations:** Chair of Phytopathology, TUM School of Life Sciences, Technical University of Munich, Emil-Ramann-Str. 2, 85354, Freising, Germany; Chair of Computational Systems Biology, University of Hamburg, Notkestrasse 9, 22607, Hamburg, Germany; Chair of Food Chemistry and Molecular Sensory Science, TUM School of Life Sciences, Technical University of Munich, Lise-Meitner-Str. 34, 85354 Freising, Germany; Computational BioMedicine lab, Institute of Mathematics and Computer Science, University of Southern Denmark, Campusvej 55, Odense, Denmark; Department for Phytopathology and Plant Protection, Institute for Phytopathology, Kiel University, Hermann Rodewald Str 9, 24118 Kiel, Germany

**Keywords:** Diversity, Early Immune Response, Ethylene, Laminarin, Phytohormones, *Phytopthora infestans*, Reactive Oxygen Species, Resistance, *Solanum chilense*, Tomato

## Abstract

Natural plant populations are polymorphic and show intraspecific variation in resistance properties against pathogens. The activation of the underlying defence responses can depend on variation in perception of pathogen-associated molecular patterns or elicitors. To dissect such variation, we evaluated the responses induced by laminarin, (a glucan, representing an elicitor from oomycetes) in the wild tomato species *Solanum chilense* and correlated this to observed infection frequencies of *Phytophthora infestans*.

We measured reactive oxygen species burst and levels of diverse phytohormones upon elicitation in 83 plants originating from nine populations. We found high diversity in basal and elicitor-induced levels of each component. Further we generated linear models to explain the observed infection frequency of *P. infestans*. The effect of individual components differed dependent on the geographical origin of the plants. We found that the resistance in the southern coastal region, but not in the other regions is directly correlated to ethylene responses and confirmed this positive correlation using ethylene inhibition assays.

Our findings reveal high diversity in the strength of defence responses within a species and the involvement of different components with a quantitatively different contribution of individual components to resistance in geographically separated populations of a wild plant species.

**Highlight:** Large-scale screenings reveal geographically distinct intraspecific differences in the dominant physiological pathogen defence responses upon glucan elicitor treatment in a wild tomato species.

## Introduction

Plant defence responses against pathogens in natural populations are often polymorphic (Kahlon and Stam, 2021a). The resistance of the host can result from the major resistance proteins or from polygenic defence mechanisms against the pathogen (Vanderplank, 1963). The later type of resistance mechanism is often considered as a basal defence, whereas major gene-mediated resistance is observed as pathogen genotype dependent. Genes encoding nucleotide-binding domain leucine-rich repeat-containing proteins (NLR) are one example of major resistance genes. *NLR* genes have been studied in natural plant populations and are reported to be diverse at the genetic level (Stam *et al*., 2019a, Van de Weyer *et al*., 2019; Witek *et al*., 2021). Similarly, members of the receptor-like proteins (RLPs) family show intraspecific variation in the presence-absence of defence responses or variability in expression patterns of these genes (Van der Hoorn *et al*., 2001; Kruijt *et al*., 2005; Kahlon *et al*., 2020, Steidele and Stam, 2020). In many cases, major gene mediated resistance is complete. By contrast, basal resistance is defined as quantitative and pathogen race-non-specific (Vanderplank, 1963). This might partially be explained by its polygenic nature, and because underlying defence reactions can be activated upon exposure to elicitors or conserved pathogen-associated molecular patterns (PAMPs). Flagellin peptides (flg22, flgII-28) and chitin have been the dominant PAMPs for studies on resistance mechanisms in plants against pathogens from bacterial and fungal lineages respectively. Many additional PAMPs have been identified e.g.VmE02 homologs, produced by various fungi and oomycetes, can trigger immunity response in *N. benthamiana* (Nie *et al*., 2018) and peptide elicitor fractions from several *Fusarium* spp. activate basal defence mechanisms in *Arabidopsis thaliana* (Coleman *et al*., 2021). One other example is laminarin, which is perceived in different plant species like *Nicotiana tabacum* (Klarzynski *et al*., 2000; Ménard *et al*., 2004), Grapevine, *Vitis vinifera* (Aziz *et al*., 2003), *A. thaliana* (Ménard *et al*., 2004), tea, *Camellia sinensis* (Xin *et al*., 2019), *Nicotiana benthamiana, Hordeum vulgare, Brachypodium distachyon* (Wanke *et al*., 2020) and Olive, *Olea europaea* (Tziros *et al*., 2021). Laminarin is an oligomeric β-1,3-glucan with β-1,6-glucan branches. β-1,3 and β-1,6-glucan are the major components of the oomycete cell wall (Aronson *et al*., 1967) and may induce defence responses similar to those provoked by elicitors from the oomycete lineage.

Several molecular mechanisms have been shown to play an important role in basal defence responses. Many of these responses happen shortly upon contact with the pathogen. In some plant-pathogen interactions basal immune responses can be quantified within the first minutes of the interaction with the pathogen or pathogen-specific molecules by measuring reactive oxygen species (ROS) production (Torres *et al*., 2006). The fine-tuning in the production of ROS is one important cue toward activating resistance mechanisms which lead to the production of various phytohormones or activation of downstream defence regulators (Ramirez-Prado *et al*., 2018). Roberts *et al*. (2019) showed the amount of ROS production varied when different tomato (*Solanum lycopersicum*) accessions were treated with flagellin peptides (flg22, flgII-28). Besides the production of ROS, phytohormones present or induced in the plant can greatly influence the resistance outcome. A higher level of salicylic acid (SA) is important in activating defence response in cultivated tomato leaves against *P. infestans* (Jeun *et al*., 2000). Genes involved in ethylene (ET) and SA pathways are important in *N. benthiamana* after infection with *P. infestans* (Takemoto *et al*., 2018). A study on potato shows that upon infection with *P. infestans* large sets of genes are upregulated at multiple time points post-inoculation. These included key marker genes involved in the jasmonic acid (JA) acid signalling pathway and genes involved in primary and secondary metabolite pathways (Tian *et al*., 2006). In cultivated tomato the negative role of abscisic acid (ABA) in resistance against *Botrytis cinerea* is regulated by repressing SA signalling (Audenaert *et al*., 2002). Whereas resistance against *Alternaria solani* is enhanced upon exogenous ABA application through defence-related gene activation and defense-related enzymatic activity of phenylalanine ammonia-lyase (PAL), polyphenol oxidase (POD) and peroxidase (PPO) (Song *et al*., 2011). Exogenous application of indoleacetic acid (IAA) in the soil resulted in *Fusarium oxysporum lycopersici* disease suppression in tomato plants (Sharaf and Farang, 2004).

The different components of the phytohormone signalling pathways can have positive or negative feedback effects on each other and thus form a complex interactive signalling network (Pieterse *et al*., 2009). These complex network topologies generated an hypothesis that one or multiple components involved in the resistance need to pass a certain threshold, in order for defence to be functional (Windram and Denbi 2015). Which factors are dominant might differ dependent on the origin of a plant or population and the pathogen in question. Such differences in dominant effective defence responses in different *A. thaliana* accessions are shown by Velásquez *et al*. (2017) against the bacterial pathogen *Pseudomonas syringae* pv. tomato (*Pst*) DC3000. They found that *Pseudomonas syringae* pv. tomato *(Pst)* DC3000 resistance was shown to be mainly mediated by an increased level of the phytohormone salicylic acid (SA) in three accessions, whereas other mechanisms were dominant in the other resistant accessions. Another study showed that between six *A. thaliana* accessions large variation in JA and SA-associated basal resistance resulted in varied resistance against the nectrotrophic pathogen *Plectosphaerella cucumerine* and the hemibiotrophic bacterium *P. syringae* respectively (Ahmad *et al*., 2011).

We used a wild tomato species *Solanum chilense* to elucidate molecular cues behind the diversity in resistance against the oomycete, *P. infestans* (Stam *et al*., 2017; Kahlon *et al*., 2021). *S. chilense* is a suitable organism to study the variation of molecular responses associated with basal defence mechanisms. Populations of *S. chilense* are geographically structured in four distinct groups based on genomic studies (Böndel *et al*., 2015; Stam *et al*., 2019a). The two southern groups are recent expansions of the species, are genetically more divergent and might be developing into new subspecies (Raduski and Igic, 2021). Thus the system provides a strong genetic structure. Previously, we found variation in defence responses against the apoplastic fungal leaf pathogen *Cladosporium fulvum* (syn. *Fulvia fulva, Passalora fulva*), with complete loss of pathogenic protein recognition in plants from the southern groups (Kahlon *et al*., 2020). In order to assess phenotypic variation in resistance, we also quantified the number of successful infection events in *S. chilense* plants after drop inoculation with various other leaf pathogens. We showed clear differences in the frequency of successful infections after inoculation of three common filamentous pathogens in different *S. chilense* populations (Stam *et al*., 2017). In a recent report (Kahlon *et al*., 2021) we showed that differences in *P. infestans* resistance between the geographically distinct populations of *S. chilense* are predominantly driven by the host genotype and can likely be attributed to differences in basal resistance, rather than isolate-specific resistance.

Here, we aim to dissect the various possible immune responses in *S. chilense* populations. We specifically investigate the early basal defence responses in *S. chilense* upon challenge with the nonspecific elicitor laminarin. We confirm that laminarin elicits a subset of defence responses triggered by *P. infestans*, we report high diversity in several key regulators of basal immune responses in *S. chilense* within and between populations of the species within the first hours of infection, and we assess their individual and joint effect on the interaction outcome, by comparing these results to our previously published infection data (Kahlon *et al*., 2021).

## Materials and methods

### Plants material used and maintained

We used 83 plants of *S. chilense* originating from nine populations (accessions) (8-10 plants each): LA1958, LA1963, LA2747, LA2932, LA3111, LA3786, LA4107, LA4117A and LA4330. The seeds of these populations were procured from the C. M. Rick Tomato Genetics Resource Center of the University of California, Davis (TGRC UC-Davis, http://tgrc.ucdavis.edu/) where the populations were originally collected as random collection of seeds from the wild populations and are now maintained and propagated at the TGRC to maintain genetic diversity. Procured seeds were sown and plants were maintained in controlled greenhouse conditions (16h light and 24°C temperature at daytime and 18-20°C at night) at TUM’s plant technology center. Each plant used in this study was at least a year old. Plants were maintained throughout the experiments by cutting them every two weeks. Each population used in this study represents one of four geographical locations of the species habitat and were originally collected during different years from wild populations. Each individual plant within a population is genetically unique.

### Evaluation of laminarin potential to activate early immune responses similar to *P. infestans* using 3’ RNAseq

We selected the central population LA3111 to evaluate differentially expressed genes in the transcriptome upon challenging with *P. infestans* Pi100 (3000 sporongia/ml) and laminarin (1mg/ml, Sigma-Aldrich) treatment (using spray inoculation). We use LA3111 population because the reference genome of *S. chilense* was generated from an individual from this population (Stam *et al*., 2019b). To measure the general defence responses in the population, the experiment was done on nine plants and all plants were pooled per treatment for the RNA extraction. Detached leaves were kept upside down in plastic boxes containing wet tissue beds and treated with water, laminarin or *P. infestans*. The boxes were kept at 18-20°C for 3 hours and samples were taken and snap-frozen in liquid nitrogen. RNA was isolated using Qiagen®RNeasy plant mini kit according to the instruction manual. Each treatment consisted of pooled samples of nine plants and four technical replicates of each treatment.

3′RNA libraries were prepared according to the manufacturer’s protocol using the QuantSeq 3′mRNA-Seq Library Prep Kit (Lexogen, Vienna, Austria). Sequencing was performed on a HiSeq2500 (Illumina, San Diego, CA, USA) with single-end 100bp reads using Rapid SBS v2 chemistry. The raw sequencing reads in FastQ format were trimmed to remove adapter sequences using Trimmomatic v0.39 (Bolger *et al*., 2014). The reads were quality filtered and trimmed using the following settings: LEADING:3, SLIDINGWINDOW:4:15, MINLEN:40. HISAT2 (Kim *et al*., 2015) was used to align sequencing reads to a reference genome (Stam *et al*., 2019b). After alignment, featureCounts (Liao *et al*., 2014) was used to identify the number of reads that mapped to genes. For feature Counts, all entries tagged as ‘gene’ were extracted from the gff annotation files and by adjusting the gene_id and transcript_id identifiers, this gene list was converted into a gtf annotation file. The downstream region of every gene was extended by 1kb (the extension stops when it hits the next gene start site). FeatureCounts was modified to search for the tag ‘gene’ instead of the default ‘exon’ tag. Differential gene expression analysis was carried out using the R package DESeq2 (Love *et al*., 2014). DESeq2 uses the output of featureCounts to estimate the fold change in gene expression between different treatment groups. Default parameters from the DESeq2 package were applied and differentially expressed genes (DEGs) showing adjusted *p*-value<0.05 were considered significant.

Gene Ontology (GO) enrichment analysis was based on previously annotated ontologies (Stam *et al*., 2019b). GO terms were selected for all candidate genes. The Background frequency of each GO term is the number of genes annotated to this GO term in all genes, while sample frequency is the number of genes annotated to that GO term in the list of DEGs in this sample.

For the manual inspection of the functions of the differentially expressed overlapping gene candidates in laminarin and *P. infestans* treated samples, we performed a BLAST search, extracted gene names and functional annotation from the best hits and if needed, performed a literature search for papers that described the functions of the described gene candidates. (See 10.5281/zenodo.5101308)

### Gene expression analysis of the key indicators of phytohormones pathways via qPCR

To independently evaluate phytohormone regulation in response to laminarin (1mg/ml) elicitation we tested the expression level of key indicators of three well-known defence phytohormones *1-aminocyclopropane-1-carboxylic acid synthases 2, ACS2* from the ET pathway (Gravino *et al*., 2015), *isochorismate synthase, ICS* (Di *et al*., 2017) and *phenylalanine ammonia-lyase, PAL* (Peng *et al*., 2004) from the SA pathway and *lipoxygenase D, LOXD;* (Heitz *et al*., 1997) from the JA pathway, in individual plant leaf discs of *S. chilense* upon treatment with laminarin and compared it to mock-treated (water) leaf discs. *S. chilense* reference names of the genes are provided in Table S1.

Leaf discs of the plant LA1963-02 (chosen due to its high resistance observed in Kahlon *et al*., 2021) were treated with laminarin and MilliQ-H_2_0 treated leaf discs served as control. Experiments were performed on three different dates in three independent replicates each.

Samples were treated for 1.5 hours, snap-frozen in liquid nitrogen and ground to a fine powder with a mortar and pestle. RNA was extracted using the Qiagen®RNeasy plant mini kit according to the instruction manual. cDNA synthesis was performed using a Qiagen QunatiTect^®^ reverse transcriptase kit according to the instruction manual. Quantitative PCR (qPCR) was performed on the synthesised cDNA using Takyon^™^ Low ROX SYBR®master mix ddTTP blue (Eurogentec Liège, Belgium). qPCR was performed in three technical replicates and a non-template control was included. Primer pairs for each tested gene are indicated in Table S1, *TIP-41* (Fisher *et al*., 2013; Nosenko *et al*., 2016) was used as a housekeeping gene for normalization and primer efficiency was performed for all the primer pairs and are shown in Table S1. The PCR reaction comprised of 10μl SYBR Green-ROX Mix, 0.3μM of forward primer and 0.3μM of reverse primer, 3μl of cDNA and volume was adjusted to 20μl with MilliQ-H_2_0. The thermal cycling profile was set to a hot start at 95°C for 3 minutes, followed by 40 cycles of amplification (95°C for 30 seconds, 60°C for 30 seconds, 72°C for 1 minute), 1 cycle melting (95°C for 30 seconds, 65°C for 30 seconds, 95°C for 30 seconds), and in the end 1 cycle at 72°C for 10 minutes. Melting curve temperatures were recorded at the end of the cycle for quality control. Data were evaluated with the software Agilent Aria MX 1.7 and relative gene expression was calculated based on the Livak and Schmittgen method (2001).

### ROS production measurement

To measure ROS production upon elicitation with laminarin, we performed a 96-well plate assay based on chemiluminescence as described by Kahlon and Stam (2021b). Leaf discs from leaves of mature plants were made with a biopsy punch (4mm, KAI Medical Solingen, Germany) and incubated in white 96-well flat-bottom plate overnight in 200μl of 20mM MOPS (pH 7.5) at room temperature. The next day buffer was removed and wells were supplemented with 75μl HRP mix (10μM horseradish peroxidase (HRP) and 10μM L012). A baseline reading was performed for the initial 10 minutes using a Luminoskan Ascent (Thermo Scientific) and then laminarin (1mg/ml final concentration) dissolved in MOPS was added to the plate at 4 wells/plant. For each population, the assay was performed on three different dates with four technical replicates each for treatment and mock (MOPS) per date. In addition, we performed ROS production measurements with flg22 at a final concentration of 500nM in population LA4330 (7 plants) to compare it with specificity of laminarin in ROS production. Normalization of data was performed by first averaging over the initial 6-10 minutes baseline reading, followed by normalization to the mock treatment for the treated leaf discs.

### Phytohormone measurements: ET measurements

Leaf discs were obtained with a 4mm biopsy punch and incubated overnight in Petri dishes containing milliQ-H_2_O at room temperature. Following overnight incubation, three leaf discs were added to glass vials (5ml) containing 300μl of milliQ-H_2_O. Laminarin was added in a final concentration of 1mg/ml to 3 glass vials (samples) containing three leaf discs of one plant and milliQ-H_2_O in three separate glass vials containing leaf discs for same plant served as a negative control. Upon addition of elicitor or water, the glass vials were sealed with septa (Carl Roth GmbH). Samples were incubated for three hours at a shaker at ~ 20-50 rpm (Heidolph Polymax 2040). 3 hours post-incubation, 1ml of air was retrieved from each samples with a syringe through the rubber cap and injected into a Varian 3300 gas chromatography machine containing AlO_3_ column with length 1m and 225°C detector temperature and 80°C column and injector temperature. The gases used for the separation of ET from the sample were H_2_, N_2_ and O_2_ at 0.5 MPa each. The amount of ET was calculated based on the standard calculation as developed by Von Kruedener *et al*. (1994) using the area under the curve (AUC). In total, we measured up to nine samples per plant on three different dates, each date containing up to three samples.

### Phytohormone and their derivatives measurements: Measurement of SA, JA, ABA, IAA, phaseic acid (PA) and dihydrophaseic acid (DPA)

Samples for measurements of these six compounds were also prepared based on the leaf disc treatment method. 150-200 leaf discs were made per plant using a 4mm diameter biopsy punch and incubated overnight in Petri dishes containing milliQ-H_2_0. The next day for each plant a 6-well plate filled with milliQ-H_2_0 was prepared for elicitation containing 25-30 leaf discs per well. Three wells were elicited with laminarin (1mg/ml) and in the remaining three wells milliQ-H_2_0 was added as a control. The plates were incubated for three hours at a shaker at ~ 20-50 rpm. Following the treatments, the leaf discs were transferred to 2ml Eppendorf tubes and residual water was pipetted out before snap-freezing the samples in liquid nitrogen.

Fine powder from the plant material was obtained after grinding the frozen leaf discs with mortar and pestle in liquid nitrogen. The samples were then processed for extraction of the phytohormones and their derivatives as described by Chaudhary *et al*. (2020), with minor modifications. 50-200mg ground material was transferred to a 2ml bead beater tube (CKMix-2ml, Bertin Technologies, Montigny-le-Bretonneux, France). 20μl of internal standard solution containing indoleacetic acid-d2 (Sigma Aldrich, Steinheim, Germany) (2.5μg/ml), salicylic acid-d4 (Olchemim, Olomouc, Czech Republic) (2.5μg/ml), (+) cis, trans-abscisic acid-d6 (Sigma Aldrich, Steinheim, Germany) (2.5μg/ml), and (-) trans-jasmonic acid-d5 (25μg/ml) (Santa Cruz, Dallas, TX, USA) were dissolved in acetonitrile and added to the samples and incubated for 30 minutes at room temperature. Following that 1ml of ice-cold ethyl acetate (Art. 864, Merck, Darmstadt, Germany) was added to the samples and stored overnight at −20°C. The next day samples were shaken for 3×20 seconds using the bead beater (Precellys Homogenizer, Bertin Technologies, Montigny-le-Bretonneux, France) at 6000rpm with 40 seconds breaks in-between. The material was then filtered with a 0.45μm pore size filter (Sartorius, Darmstadt, Germany) using a Minisart^®^ syringe. The filtrate was transferred to 2ml tubes and vacuum dried. Then samples were reconstituted in 70μl of acetonitrile and sonicated for 3 minutes. 2μl of the sample from the HPLC tubes (glass vials) were injected into the LC-MS/MS system. The MS method used measured positive and negative ionization mode within one run (polarity switching). Negative ions were detected at an ion spray voltage of −4500 V (ESI-) using ion source parameters: curtain gas (35 psi), temperature (550°C), gas 1 (55 psi), gas 2 (65 psi), collision activated dissociation (−3 V), and entrance potential (−10 V). Positive ions were detected at an ion spray voltage at 4500 V (ESI+) using ion source parameters: curtain gas (35 psi), temperature (550°C), gas 1 (55 psi), gas 2 (65 psi), collision activated dissociation (−3 V) and entrance potential (10 V) and 40°C column oven temperature was at a QTRAP 6500+ mass spectrometer (Sciex, Darmstadt, Germany). MS/MS fragmentation was obtained and samples were separated by ExionLC UHPLC (Shimadzu Europa GmbH, Duisburg, Germany) using 100 × 2.1 mm2, 100 Å, 1.7 μm, Kinetex F5 column (Phenomenex, Aschaffenburg, Germany). Solvent used for separation were (A) 0.1% formic acid in water (v/v) and (B) 0.1% formic acid in acetonitrile (v/v) with a flow rate of 0.4 ml/minute. Chromatographic separation was performed with the gradient of 0% B for 2 minutes, increased in 1 minute to 30% B and in 12 minutes to 30% B, increased in 0.5 minute to 100% B, held 2 minutes isocratically at 100% B, decreased in 0.5 minute to 0% B, and held 3 minutes at 0% B. Phytohormone quantification was performed based on comparison with standard curves prepared with purified hormones and using AUC. The final concentrations were obtained in nanograms of hormone per gram of fresh weight of the sample.

### Infection data

Data on *P. infestans* infections were taken from Kahlon *et al*. (2021), using the same methods that were previously described in Stam *et al*. (2017). In these studies, detached leaves were drop infected with a *P. infestans* solution (3000 sprongia/ml). All leaflets of the compound *S. chilense* leaves were infected with a single drop and the infection frequency (IF) was calculated per leaf and summarized per plant an population. The data originate from the exact same plants as those used in this study.

### Statistical analysis of the data, Pearson’s correlation and linear mixed models for infection frequencies with components of basal immunity and stress-related phytohormones

All the data analyses were performed in the R software (version 3.4.4, R core Team, 2020). ANOVA was performed with the function aov(), and post hoc Tukey tests with the function TukeyHSD(), from the package {stats}. When the *p*-value was lower than 0.05 it was considered significant. Pearson’s correlation was performed using the function cor(). The analyses were done for the 83 plants for which IF scores were available (Kahlon *et al*., 2021). Figures were made using the R package {ggplot2}.

### Validation of ET accumulation in delivering resistance in individuals from the southern coastal population

ET validation experiments were performed on two individuals (plant 05 and plant 10) from a southern coast population, LA4107, selected based on Pearson’s correlation among ET production and infection frequency. ET measurement in the leaf discs was performed as described above. ET blocking was performed by adding 5μM aminoethoxyvinyl gylcine (AVG) (Sigma) to the samples and 3 hours post-treatment ET was measured by gas chromatography. The infection frequency upon treatment with 5μM AVG was determined using detached leaf infection assay as described in Kahlon *et al*. (2021): detached leaves were surface sterilized with 70% ethanol and kept upside down (adaxial side facing upwards) in plastic boxes containing a wet tissue bed. The leaf set for AVG treatment was kept on a wet tissue bed made with water containing AVG (5μM) and leaves were sprayed with 5μM AVG following drop inoculation with *P. infestans* isolate Pi100 (3000 sporangia/ml). The experiment was repeated on eight individual leaves per treatment. Throughout the experiment, 18-20°C temperature was maintained and boxes were kept in dark. The infection outcome was taken at 7 days post-inoculation.

## Results

### Laminarin elicits a transcriptional response overlapping with that induced by *P. infestans*

First, we set out to confirm whether the purified glucan elicitor laminarin can elicit oomycete-like early defence responses in *S. chilense*. Therefore, we infected *S. chilense* plants of population LA3111 with *P. infestans* or treated the plants with laminarin and measured the transcriptional response after 3 hours.

As expected, infection with *P. infestans* triggered strong transcriptional responses. In total, we measured 595 differentially expressed genes (false discovery rate adjusted *p*-value<0.05) upon *P. infestans* infection (371 up-regulated and 224 down-regulated). Laminarin treatment results in 102 differentially regulated genes (31 up-regulated and 72 down-regulated). (Table S2). More than 50% of the genes that were differentially expressed after laminarin treatment were overlapping with the *P. infestans*-associated response. For the upregulated gene fraction, the overlap was 77%. Only a small number of DEGs (12) can be uniquely detected in a direct pairwise comparison between the Laminarin-treated and *Phytophthora*-treated samples, thus the false positive discovery rate in this experiment is likely lower than 2%. (Figure 1A-C). To validate our hypothesis that laminarin triggers a decomplexified defence response, we analysed the annotations of the overlapping gene lists. 35% of the overlapping genes are associated with the Biological process: Stress Response and the overlapping fraction is significantly more often annotated with the GO terms enzyme regulator activity and receptor activity (chi-square test, p < 0.01, Figure 1D-E, Table S3-S4). Moreover, homologs of more than half of the overlapping genes are reported in the literature to be involved in defence responses (Table S5). We found homologs of regulators of the plant ROS response (SOLCI005830700, Peroxidase *CEVI1*), key regulators of defence hormone signalling like *ER5* (SOLCI004643500, ET response); *LOX1* (SOLCI003764800, JA signalling) or *PAL3* (SOLCI000597200, SA signalling) and upregulation of *Mitogen-activated protein kinases* (*MAPKs*, SOLCI002491100). This supported that laminarin can trigger a subset of oomycete-associated defence responses and is a suitable compound to study variation in basal defence responses in *S. chilense*.

**Figure 1:**
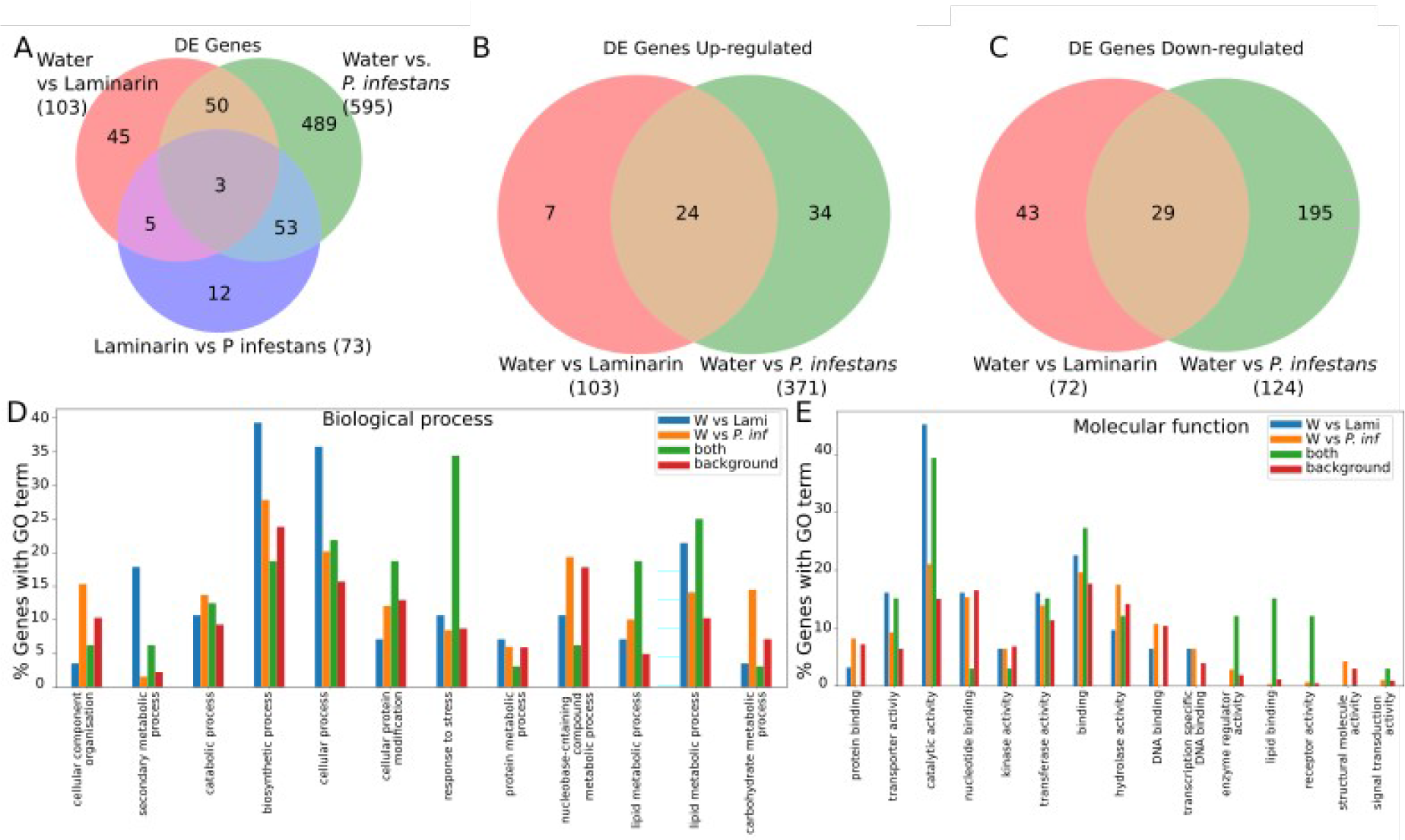
RNAseq analysis of the central population LA3111 (nine plants pooled per treatment) of *S. chilense* 3 hours after *P. infestans* (3000 sporongia/ml), laminarin (1mg/ml) and water treatment. Differentially expressed genes (DEGs) overlap in different treatment: overall (a), up-regulated (b) and down-regulated (c), Gene Ontology (GO) analysis of RNA-seq data showing % genes with signals for gene ontology terms for biological and molecular function for laminarin Vs *P.infestans* treatment (d,e).

To independently verify the involvement of laminarin-specific phytohormone-associated defence responses, we evaluated the expression of key regulators described in literature of three phytohormones, ET, SA and JA in an *S. chilense* individual from a different central population: LA1963 (plant 02). We observed an up to 7.8-fold increase in expression of *S. chilense ACS2* (SOLCI000989600), a key regular in the ET pathway in laminarin treated samples as compared to water-treated samples (Figure S1). Next, we looked into key regulators from the two known pathways for SA regulation. We observed that *PAL*-like transcripts (SOLCI002546900) showed up to 14-fold increase in laminarin-treated samples as compared to water-treated controls. *ICS* (SOLCI004470400), the key regulator of the second SA pathway was downregulated (Figure S1). For the JA pathway, we performed qPCR on the *LOXD* (SOLCI003768300) gene and observed an up to 10-fold increase in the laminarin treated samples (Figure S1). Thus, we confirmed differential regulation of key regulators in defence-associated phytohormone pathways.

### ROS production in *S. chilense* is highly polymorphic

ROS is one of the important key regulators in basal immune responses, and regulators of the ROS pathway were differentially expressed in the RNAseq data (Table S5). Thus, we tested ROS production in 83 genetically distinct *S. chilense* plants upon elicitation with laminarin. Maximum ROS production upon laminarin elicitation was significantly different between the populations (Figure 2A, Table S6). The highest average ROS maximum was recorded in LA3111 and the lowest average in LA3786.

**Figure 2:**
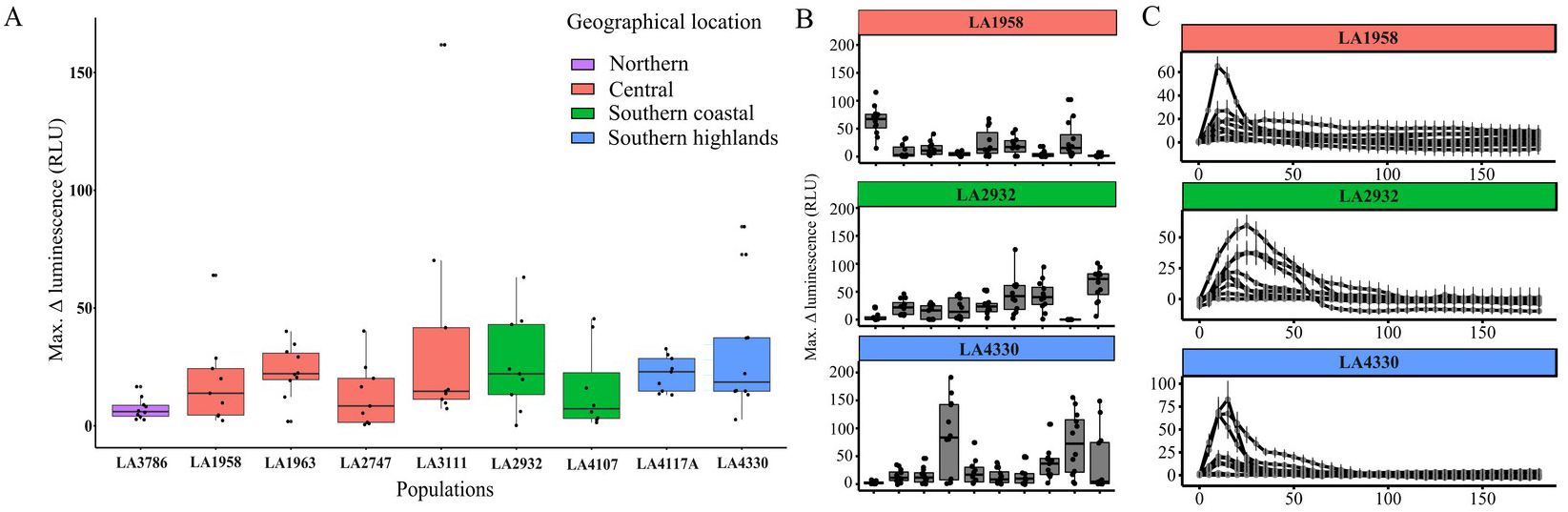
ROS accumulation in the leaf discs from *Solanum chilense* measured from 0-180 minutes upon elicitation with laminarin (1mg/ml). A) Overview for each of the populations. Each boxplot represents a populations, each black dit the mean measured value for one plant, obtained from three or four individual repetitions, as explained in B, All the significance data is highlighted in supplemental material. B) examples highlighting the within-population diversity for for three of nine populations. Each box plot represents an individual plant per population with data from one leaf disc represented as one data point accounting up to ten to twelve leaf discs per plant. Individual measurements were performed on three different dates (n=3-4 each date; 3×3(4)=10(12) leaf discs per plant), each median of the boxplot B represnt single datapoints of A in corresponding population plant datapoints. C. Differences in ROS kinetics for the three highlighted populations from panel B. X-axis are minutes post treatment. Y-axis shows relative luminescence unit (RLU). Colors of the boxplots or header bars represent the geographical location of the population. Extended panels like B and C for all populations can be found in Figure S3 and S4. Each data poiunt is a median similar to panel B and the error bar represnet standard error to have a clear visulatization of the different plants.

When grouping the populations in geographical region and looking at the overall average in ROS maxima the southern highlands group had the highest ROS production and the northern group had the lowest (Figure S2). We also found significant differences in ROS maxima in within the individual populations for 8 out of 9 populations (Table S7), with some plants showing high ROS production and others showing no detectable ROS production upon elicitation with laminarin (Figure 2B and Figure S3).

We also observed variation in the kinetics of the ROS production, with plants in some populations not showing a clear single peak, but rather a longer-lasting ROS production. This phenomenon appeared more common in the southern populations and occurs most frequently in southern populations (Figure 2C and S4).

To confirm the specificity of the observed ROS production towards laminarin, and to show that lack of observed ROS burst does not result from a general ROS signalling impairment in the plants, we further tested ROS production after elicitation with the bacterial PAMP peptide flg22 in all plants from population LA4330 (Figure S5). This revealed variable ROS production upon elicitation with flg22. Moreover, there appears to be no apparent correlation between the strengths of flg22- and laminarin-triggered responses. Some plants showed no flg22 response and a clear laminarin response, or vice versa and some plants showed responses in similar intensity. This suggests that the observed differences in some plants are elicitor-specific variation and not a general effect of ROS production ability or defence signalling pathways.

### ET accumulation upon laminarin treatment is low in southern highland populations

To evaluate the role of phytohormones in the resistance differences observed in *S. chilense*, we first looked into ET production. Upon elicitation with laminarin, we observed that plants showed significant differences in ET production as compared to mock-treated samples (Figure 3, Table S8 and S9). Out of 83 plants tested, we found significantly induced ET production upon elicitation in 39 plants. Plant 02 from LA1963 showed a clear ET response and so do several plants from population LA3111, confirming our RNAseq and gene expression qPCR results above (Figure S1 and Table S9). Hence, differential ET-pathway gene expression in these plants leads to laminarin-elicited ET accumulation.

**Figure 3:**
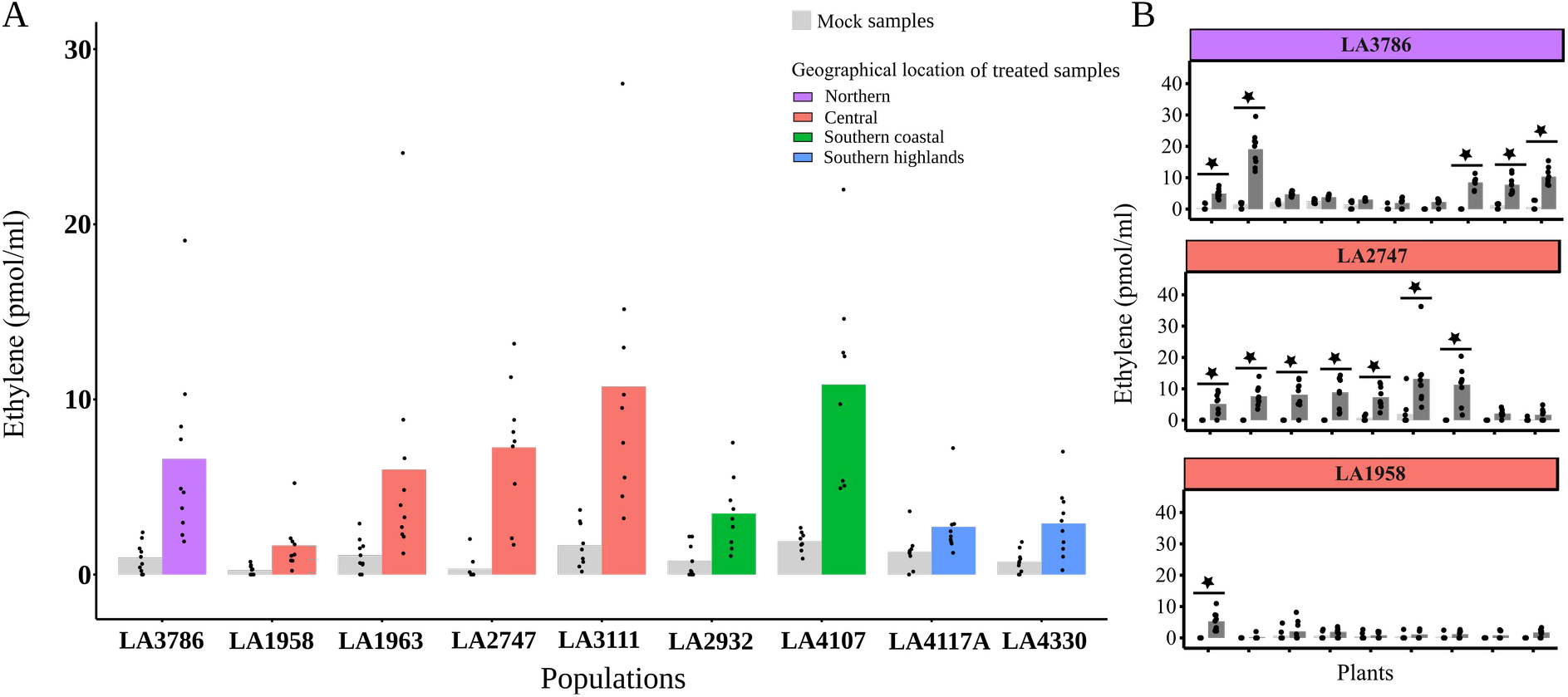
ET accumulation in the leaf discs from *S. chilense* 3 hours upon elicitation with laminarin (1mg/ml) and mock (milliQ H_2_0). A) Each bar pair, (light grey and colored) represents an population. Each bar schows the mean of the population each dot represents the mean of one plant from three individual repetitions (as in B), All the significance data is highlighted in supplemental material. B) Each bar pair (light grey and dark grey) represents an individual from the population. Each bar shows mean of 7-9 data points which represent 7-9 samples measurements performed on three different dates (n=2-3 samples each date), each sample contained three leaf discs. Significantly different ET accumulation in laminarin treated samples from the mock treated samples in an individual is represented with the star on the bar pair (ANOVA with post hoc Tukey tests on complete dataset). Y-axis shows ET accumulation in pmol/ml headspace of the samples. Each panel in B shows different population and colors represent the geographical location of the population. Panels for all additional populations can be found in Figure S6A.

We find significant differences in ET accumulation between the populations (Figure 3A, Table S10). Looking at the populations based on their geographical locations shows that overall average of ET accumulation was lowest in the southern highlands group (Figure S2B). Within populations, we observed that the number of plants that significantly differ in ET response varied dependent on the population. Population LA3786 showed the most differences between individual plants and LA2747 was the most uniform, population LA1958 showed nearly no ET response (Figure 3B, Table S11).

### Populations show high diversity in the basal level of phytohormones

Next, we looked into the production of two important defence-related phytohormones and their derivatives, for which we detected expression of key regulators in our RNAseq data, JA and SA, as well as other phytohormones and their derivatives that are known to be involved in stress responses: ABA, PA, DPA and IAA.

With our method, we were unable to detect quantifiable amounts of JA in any of the samples tested, but were able to quantify both free and total SA in the basal state and after elicitation. We observed that most populations did not show strong differences in the levels of SA (free and total) after elicitation with laminarin (Figure 4A and B, Figure S6 B and C and Table S8). Four plants form an exception: two showed a higher amount of free SA (LA3111 plant 05 and LA4330 plant 05), and two showed a lower amount of free SA (LA2932 plant 12 and LA4107 plant 12) when compared to the mock-treated samples (Figure S6B). These differences did not correspond with those plants being more resistant or susceptible than other plants in the corresponding populations.

**Figure 4:**
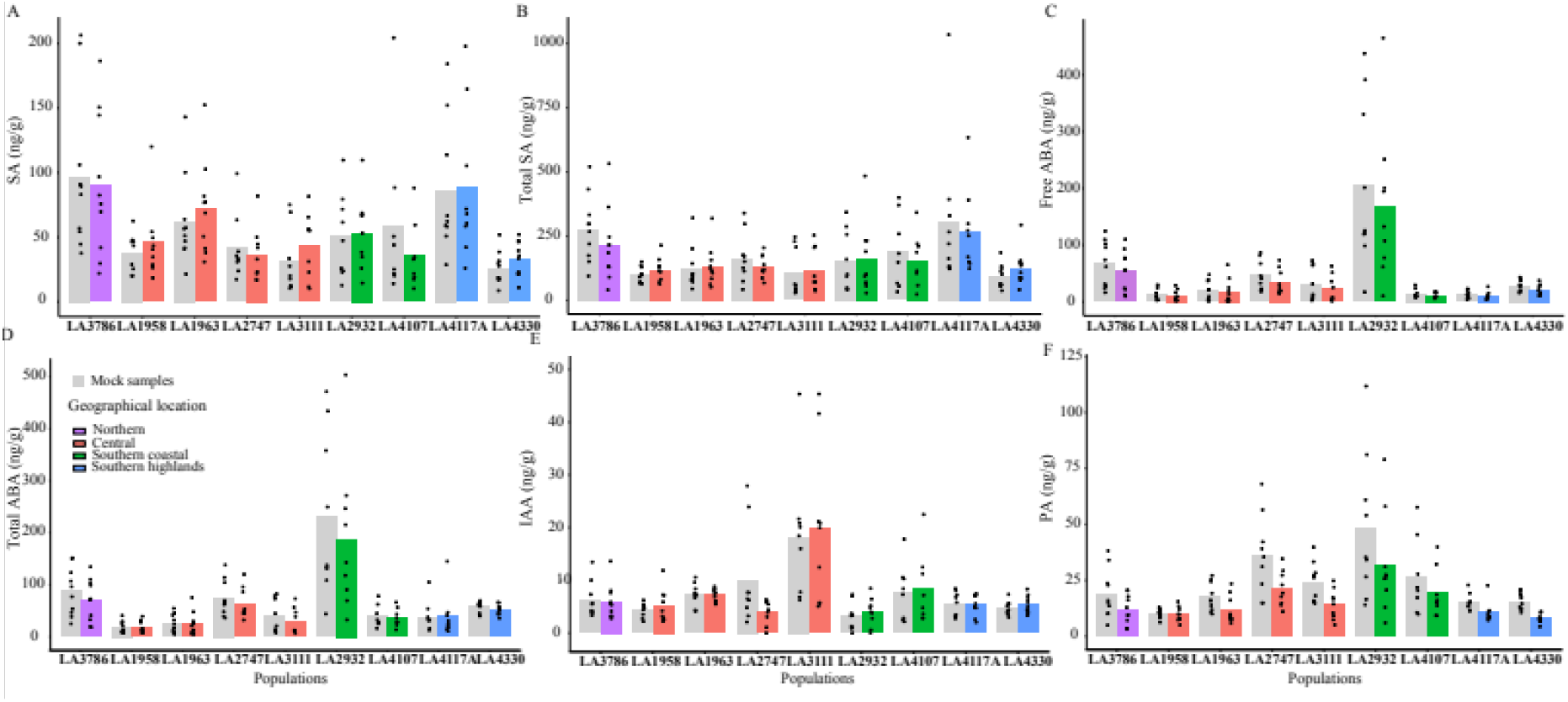
Phytohormone measures in the leaf discs from *S. chilense* 3 hours upon elicitation with laminarin (1mg/ml) and mock (milliQ H_2_0). Each bar pair (light grey and colored) represents a population. Each dot represents the mean of a single individual measured with at least three independent repetitions. Results for each individual plant can be found in Figure S6. Y-axis shows phytohormone accumulation in ng/g of the samples. Colors represent the geographical location of the population.

Interestingly, we measured significant differences in basal levels of free and total SA within and between populations. The correlation coefficient between free and total SA content is 0.46 (*p*-value=1.10E-05) (Table S12), therefore we treated free and total SA independently in further analyses. Both basal SA levels (free and total) were significantly different between the populations (Table S13). As with ROS and ET responses, we also found significant differences within the populations for basal levels of both free SA and total SA content (Table S14).

We also measured ABA, PA, DPA and IAA, which are described to be important for biotic stress responses and pathways of several of these hormones influence each other. We did not detect DPA in our samples. We detected basal levels for the phytohormones ABA (free ABA, Figure 4C, Figure S6D and total ABA, Figure 4D, Figure S6E), IAA (Figure 4E, Figure S6F) and PA (Figure 4F, Figure S6G),. Laminarin treatment did not significantly change the level of these phytohormones except for PA (Table S8). For PA we observed a significantly lower amount after treatments when compared to the basal level in plants (Figure 4F, Figure S6G and Table S15). Although, in general basal levels of PA and PA levels upon elicitation were highly correlated (Pearson’s correlation coefficient of 0.74, *p*-value=2.2E-016) (Table S12). We observed a higher amount of basal levels of free and total ABA in LA2932 as compared to other populations, whereas IAA was higher in LA3111. The levels of all these phytohormones show significant differences between and within populations (Table S16 and Table S17, respectively) in an independent manner (Table S12). Looking at the data of all the tested populations based on geographic location, we found higher levels of basal PA in the southern coast and IAA was higher in the central region whereas SA (free and total) was high in the north and ABA levels were higher in northern and south coast populations (Figure S2).

### Multiple defence responses correlate with observed resistance phenotypes

To assess whether the individual defence responses measured in the plants can be associated with *S. chilense* resistance properties, we looked for correlations with previously generated data on the frequency with which *P. infestans* can infect *S. chilense* leaflets, the so-called infection frequency (IF) (Kahlon *et al*., 2021). We found no correlation between the observed ROS maxima and the IF observed with *P. infestans* (Pearson’s correlation coefficient of 0.09, *p*-value=0.37, Table 1), whereas we found a significant negative Pearson’s correlation of IF with ET accumulation (−0.36, *p*-value=0.0008; Table 1). The Pearson’s correlation of IF with basal levels of PA also showed a negative correlation (−0.2443413, *p*-value=0.026; Table 1), whereas we observed no strong or significant (*p*-value<0.05) correlation for SA, ABA and IAA (Table 1).

**Table 1:** Pearson’s correlation of measured potential immunity-related factors at basal levels and upon elicitation with laminarin (1mg/ml) with the infection frequency (IF) of same plants upon inoculation with *P. infestans* Pi100 published in Kahlon *et al*. (2021). The correlation is shown for the all the measured components. Significant correlation is highlighted in green.

| Parameter compared | Pearson's correlation coefficient | *p*-value |
| --- | --- | --- |
| IF~ROS | 0.10 | 0.3748 |
| IF~ET | -0.36 | 0.0008 |
| IF~Free SA (Basal) | -0.05 | 0.6290 |
| IF~Total SA (Basal) | -0.15 | 0.0690 |
| IF~Free IAA (Basal) | -0.13 | 0.2727 |
| IF~Free ABA (Basal) | -0.12 | 0.2737 |
| IF~Total ABA (Basal) | -0.12 | 0.2419 |
| IF~PA (Basal) | -0.24 | 0.0260 |

### The dominance of individual defence responses differs geographically

All measured components showed geographical trends at basal and induced levels (Figure S2, as well as S3, S4, S6). This supports that the plants’ genotypes rather than the common experimental environment was the driver of metabolic differences between the plants. We hypothesize that different populations have adapted different defence strategies due to adaptation to specific climatic niches. Thus, to confirm the possible larger effect any of measured components in certain geographical regions, we calculated the correlation coefficient for ET and PA for each of the geographical groups of *S. chilense*. We found that the effect of ET is most strongly correlated with resistance in the coastal populations, whereas PA showed the strongest correlation to resistance in the central group (Figure 5 and Table S18).

**Figure 5:**
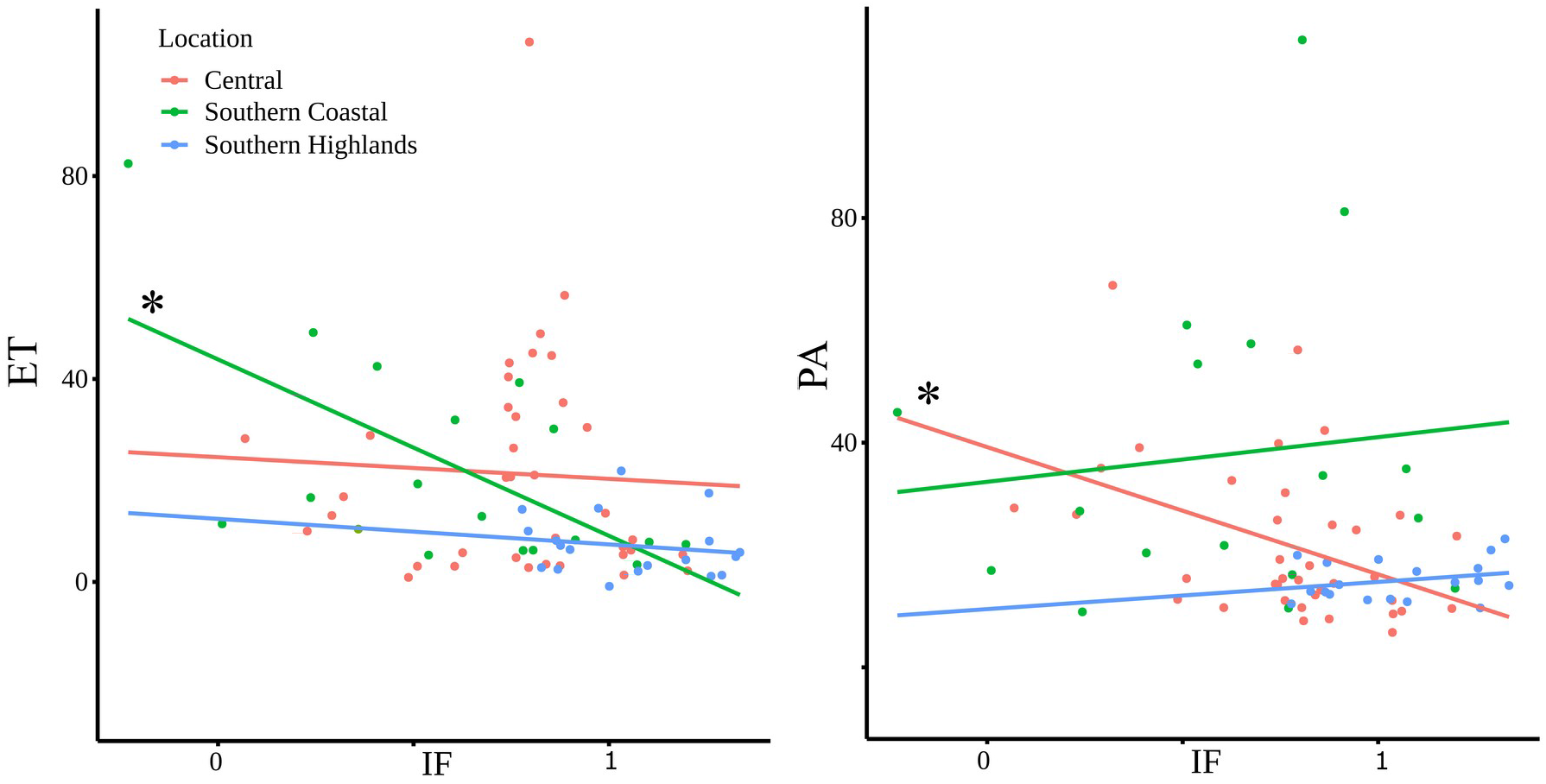
Correlations between the *P. infestans* infection frequency (IF, X-axis) and the measured ET or PA response phytohormone accumulation in ng/g of the samples or pol/ml respectively (Y-axis). Each dot represents the mean value of an individual plant from three individual repetitions. Infection frequencies were obtained from and independent experiment from the same plants as presented in Kahlon et al (2020). IF at zero indicates plants that are fully resistant and IF at one shows 100% infection rate upon inoculation. The pearson correlations were calculated per geographic group. The * indicates the significant correlations for the central and southern coastal populations for PA and ET respectively.

### ET plays a role in defence response in southern coast tested individuals

To confirm the contribution of ET to resistance in the coastal populations, we selected two plants from southern coastal population LA4107, one with relatively high ET production and another with relatively moderate ET production, with low and medium-high scores from the infection frequency spectrum, respectively. To verify the role of ET on the resistance outcome, we used AVG, a well-established chemical inhibitor, to halt the ET production in the selected plants and tested the ET accumulation after laminarin treatment. AVG was successfully able to inhibit the ET production up to 100% in the plant samples (Figure 6A). After inoculation with *P. infestans* isolate Pi100, the plants indeed showed higher susceptibility when they were pre-treated with AVG as compared to control plants, confirming the positive role of ET in basal resistance in this population (Figure 6B).

**Figure 6:**
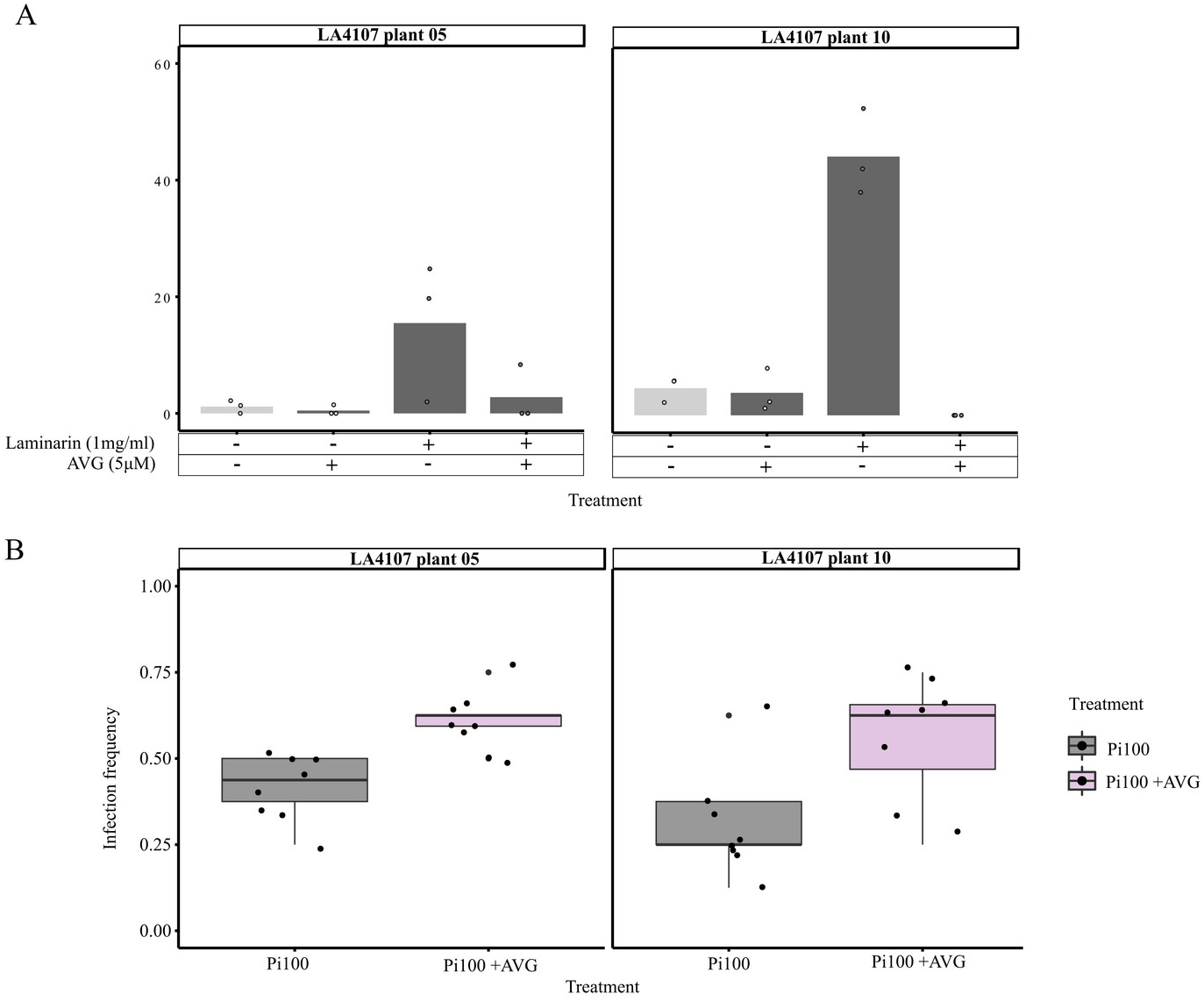
ET inhibition assay on LA4107 plant 05 and plant 10: ET accumulation in the the leaf discs from *S. chilense* LA4107 plant 05 and 10, 3 hours upon elicitation with laminarin (1mg/ml), AVG (5μM), laminarin (1mg/ml) + AVG (5μM), and mock (milliQ H_2_0). Light and dark grey crossbar pair represents plant with and without treatment with laminarin (1mg/ml). Each crossbar is the mean of three samples measurement. Y-axis shows ET accumulation in pmol/ml air of the samples. b) Detached leaf infection assay of LA4107 plant 05 and 10 upon drop inoculation with *Phytopathora infestans* Pi100 (3000 sporangia/ml) with and without AVG (5μM) treatment. Y-axis represents infection frequency which is the ratio of infected leaflets divided by inoculated leaflets. Each dot represents the ratio from one leaf, the red dot represents the mean value and the *p*-value is shown on the top of the boxplot.

## Discussion

We previously used natural populations of *S. chilense* to show intraspecific variation in resistance against *P. infestans* (Kahlon *et al*., 2021). In this study, we evaluated several key components of basal defence responses in the same plants to explore molecular cues behind the previously observed phenotypic variation.

In order to reliably and reproducibly study defence components in this polymorphic plant species, and to rule out variation arising during the preparation of pathogen biological material, we used the glucan elicitor laminarin. Laminarin has been previously reported to activate basal immune responses such as ROS production, calcium influx and MAPK activation in members of the Solanaceae family (Meénard *et al*., 2004; Wanke *et al*., 2020).

We observed significant overlaps in DEGs in *S. chilense* central population LA3111 upon elicitation with laminarin and infection with *P. infestans*. The majority of these genes are known for involvement in defence responses. We further showed differences in transcript levels via qPCR of key regulators of defence-related phytohormones after laminarin treatment in a plant of a different central population, LA1963. This suggests that laminarin can be used as a proxy for evaluating early basal immune responses activation in *S. chilense*.The RNAseq data of both *P. infestans* and laminarin treatments revealed regulation of homologs of several previously identified genes known in major defence pathways, like ROS production and both SA and JA signalling. These basal immunity components have been shown to be involved in *P. infestans* resistance in different Solanaceous plant species. In cultivated tomato, reduced accumulation of ROS results in enhanced resistance against *P. infestans* (Cui *et al*., 2016). Higher SA levels have a positive effect on *P. infestans* resistance in *S. tuberosum* (Halim *et al*., 2007). Both, SA and ET contribute to resistance in *N. benthamiana* (Shibata *et al*., 2010), and higher levels of JA and interplay with SA were observed in resistant cultivars of *Capsicum annuum* (Ueeda *et al*., 2005). We observed high intraspecific diversity in the above-mentioned components early after elicitation with laminarin, and at basal levels. Large-scale intraspecific variation in basal immunity has also been reported on a transcriptional level in *A. thaliana* accessions, upon elicitation with the bacterial PAMP flg22 (Winkelmüller *et al*., 2021).

Surprisingly, we did not find a strong correlation between the amount of ROS produced in a plant after elicitation with laminarin and its resistance properties. ROS production upon biotic stress has often been considered as a hallmark of successful recognition of pathogens and the activation of defence (Torres, 2010). ROS production linked to the perception of flg22 is often taken as an indicator for resistance against bacterial *Pseudomonas* spp. pathogens (e.g. in *A. thaliana*; Smith and Hesse, 2014 and in tomato; Roberts *et al*., 2019). Our study shows that laminarin-triggered ROS production in *S. chilense* cannot be used to estimate the basal resistance against *P. infestans*. Similarly, we also found no correlation between laminarin-triggered SA production or basal SA levels and *P. infestans* resistance. Thus, these individual defence components triggered by laminarin either have a rather limited contribution to the observed *P. infestans* resistance in the populations, or ROS and SA are not directly involved in *P. infestans* resistance in *S. chilense*. On the other hand, laminarin has been shown to induce the ET pathway, but only sulphated laminarin (ß-1,3 glucan sulfate) can induce the salicylic acid signaling pathway in *N. tabacum* and *A. thaliana* (Meénard et al. 2004). In our RNAseq analysis, we do see more DEGs when plants are treated with *P. infestans* as compared to laminarin. In the future, it would be interesting to evaluate the effects of sulphated laminarin and other known PAMPs from *P. infestans* in order to dissect the defence responses further.

Our data does support that resistance observed in our plant species can be correlated to different components in the plants: induced ET and also basal levels of the phytohormone PA. This is in line with the hypothesis that basal defence is regulated by a complex network of interacting components from plants and pathogens (Windram and Denbi 2015, Kahlon and Stam, 2021a). We also observed that the strength of the correlation is dependent on the geographical region from which the plants originated. Calculations per population would be even more interesting, but due to the limited number of plants per population, these calculations would lack statistical power.

It can be assumed that in each populations multiple components play important roles, but that due to sample size limitations, these effects were not picked up. The generation of generalized linear mixed models, testing the combined effect of multiple components would be desirable in this context, though this would require a lot of additional data.

It has previously been shown that ROS production leads to SA production in a feed-forward loop in defence responses in *Arabidopsis* (reviewed by Herrera-Vásquez *et al*., 2015). We did not observe such correlation among ROS production and SA production at early time points, nor did we observe a correlation between SA and ET production as observed in tomato resistance against the fungal pathogen *Fusarium oxysporum* (Di *et al*., 2017). Interestingly, the suppression of ABA biosynthesis and activation of ET biosynthesis upon copper ions treatment enhances resistance against *P. infestans* in potato seedlings (Liu *et al*., 2020b), whereas in our system ET positively contributed to resistance observed against *P. infestans*.

In our assays, ET had the strongest role in early defence. ET has previously been described in association with various defence responses. In *A. thaliana*, Resistance to Powdery Mildew 8 (RPW8)-mediated defence response is regulated by a ET-mediated feedback loop (Zhao *et al*., 2021). For Solaneceous species, the activation of defence-related genes in *P. infestans* resistant potato cultivars upon exogenous ET treatment has also been reported in a recent transcriptome study, by Yang *et al*., (2020) and laminarin triggers the expression of ET-dependent defence genes in *N. tabacum* (Meénard *et al*., 2004). A positive role of ET production has been reported in relation to resistance to *C. fulvum* in tomato plants carrying corresponding resistance genes against a specific *C. fulvum* race (Hammond-Kosack *et al*., 1996). Another study showed that ET is also involved in resistance to the fungal pathogen *B. cinerea* and certain wound responses in tomatoes, with no clear role of JA or SA observed (Díaz *et al*., 2002).

We further showed geographical variation in basal and induced levels of each component and expect that the role of each component might differ between populations due to variation in habitats. Interestingly, the role of ET was stronger in the coastal populations and experimentally verified with ET inhibition assays in plants from coastal population LA4107. The stronger association of ET and resistance specifically in the southern coastal populations could be a result of specific adaptation processes in these populations. This could potentially reflect an added benefit of the development of stronger ET signalling in these populations as a result of specific habitat adaptation e.g. to deal with potential abiotic stresses, like salt or temperature. General temperature dependency of defence regulation and the involvement of phytohormone signalling has been shown for both cold (Wang *et al*., 2019) and heat stress (Huang *et al*., 2021). ET has been shown to be a crucial phytohormone when it comes to coping with salinity stress in plants (Riyazuddin *et al*., 2020). The positive effects of ET in salt tolerance have been illustrated in *A. thaliana* (Yang *et al*., 2013) and *Zea mays* (Freitas *et al*., 2018). In a study by Kashyap *et al*., (2020), *S. chilense* plants under salt stress coped better than cultivated *S. lycopersicum* due to a better anti-oxidant system. We also observed high basal levels of the phytohormone ABA in the coastal population LA2932 (Figure 4C-D). ABA is highlighted to be an important phytohormone for abiotic stress tolerance including salinity stress (reviewed by Zhu, 2002 and Ng *et al*., 2014).

The genotype-to-phenotype linkages in systems biology are complex. In a diverse panel of wild and domesticated tomatoes basal resistance against the generalist fungal pathogen *Botrytis cinerea* has been reported to be dependent on the interaction of multiple loci among both host and pathogen (Soltis *et al*., 2019). Here we presented a decomplexification approach, where more components can be added to understand both the triggers (by testing different elicitors) and the outcomes (by measuring more responses). As highlighted by Marshall-Colón and Kliebenstein, (2019), such future studies should be performed using large-scale metabolomics and transcriptomics analyses, to determine the key regulators underlying the measured responses and to be able to appreciate the intrinsic value of the complexity of signalling networks.

In our previous studies (Stam *et al*., 2017; Kahlon *et al*., 2021) we showed no strong signs of host adaptation towards resistance to *P. infestans*, as resistance shows no clear geographical pattern of adaptation. However, our current results indicate that different coping mechanisms are present in each of these populations, possibly due to specific adaptation to the niches that the plants inhabit in each region. Examples of such specific adaptations have also been observed for other host-parasite interactions. Populations of *Eruca vesicaria* (syns. *Eruca sativa*, wild rocket) from Mediterranean and desert habitats showed activation in defence responses via two different mechanisms when challenged with generalist herbivore *Spodoptera littoralis*. Mediterranean plants showed accumulation of glucosinolates and desert plants showed induced levels of a specific protease inhibitor (Ogran *et al*., 2016). Beevan *et al*. (1993) showed immense differences in phenotypes of two populations of *Senecio vulgaris* (groundsel) against *Erysiphe fischeri* and proposed different populations have evolved different survival strategies against the same pathogen. In *A thaliana*, combined effects of genetic variation and differences in environmental factors also shape defence-associated metabolite contents (Katz *et al*. 2021)

Together our data support high complexity of *S. chilense’*s defence response to the general glucan elicitor laminarin. These responses might contribute to *P. infestans* resistance by additive or network functions. At single geographic location, certain plant hormones play a bigger role than at others. We hypothesize, that wild plants adapt to the local abiotic environment and hormones may be key to this adaptation. The corresponding defence machinery might simultaneously undergo co-adaptation to cope with biotic stress. We speculate that plants’ high connectivity between abiotic and biotic signaling results in the necessity to habitat-specifically recruit different defence pathways and that the nature of the involved hormones accordingly differs in the wild. This would be determined not only by the local pathogens but also strongly by the abiotic environment and highlights the need for further population-scale studies on pathogen resistance mechanisms (Kahlon and Stam 2021).

## Acknowledgments

We would like to thank Dr. Stefanie Ranf (TUM, Chair of Phytopathology), Dr. Christina Steidele (TUM, Chair of Phytopathology) and Dr. Harald Schempp (TUM, Chair of Phytopathology) for their help with setting up of ROS production assay and ET measurements assay for the wild tomato. Ethan Weiner (visiting RISE-DAAD scholar, TUM, Chair of Phytopathology) for assistance in the initial set-up of the SA and JA measurements in wild tomato, Lina Muñoz (TUM, Chair of Phytopathology) for helping with the sample preparation for phytohormones extraction, Dr. Christine Wurmser (TUM, NGS@TUM) for running the sequencing for our RNAseq experiments, Prof. Aurélien Tellier (TUM, Section of Population Genetics) for sharing the *S. chilense* populations and Sabine Zuber, Bärbel Breulmann and Anneliese Keil for maintaining them at the TUM’s plant technology center.

## Author contributions

Conceptualization: RS, PSK, CD, RHü and JB; Investigation: PSK, AF, MM, MO, MG and RHa; Data interpretation and evaluation: PSK, MM, AF, MO, RHü and RS; Writing and Data representation: PSK and RS. All authors reviewed and approved the manuscript.

## Data Statement

Analytical results are included in the supplementary data files. Raw sequence data (Illumina reads) are uploaded to NCBI SRA and available under PRJNA746795. All scripts used for the analyses, as well as all intermediate data files (e.g. FeatureCount output files) can be found on Zenodo (10.5281/zenodo.5101308)

## References

Ahmad S, Van Hulten M, Martin J, Pieterse CMJ, Van Wees SCM, Ton J. 2011. Genetic dissection of basal defence responsiveness in accessions of *Arabidopsis thaliana*. Plant, Cell & Environment 34, 1191–1206.

Audenaert K, De Meyer GB, Höfte MM. 2002. Abscisic acid determines basal susceptibility of tomato to *Botrytis cinerea* and suppresses salicylic acid-dependent signaling mechanisms. Plant Physiology 128, 491–501.

Aronson JM, Cooper BA, Fuller MS. 1967. Glucans of Oomycete Cell Walls. Science 155, 332–335.

Aziz A, Poinssot B, Daire X, Adrian M, Bézier A, Lambert B, Joubert J-M, Pugin A. 2003. Laminarin elicits defense responses in grapevine and induces protection against *Botrytis cinerea* and *Plasmopara viticola*. Molecular Plant-Microbe Interactions 16, 1118–1128.

Barragan AC, Weigel D. 2021 Plant NLR diversity: the known unknowns of pan-NLRomes. The Plant Cell 33, 814–831.

Bates D, Mächler M, Bolker B, Walker S. 2015. Fitting Linear Mixed-Effects Models Using lme4. Journal of Statistical Software 67, 1–48.

Bevan JR, Clarke DD, Crute IR. 1993. Resistance to *Erysiphe fischeri* in two populations of *Senecio vulgaris*. Plant Pathology 42, 636–646.

Bolger AM, Lohse M, Usadel B. 2014. Trimmomatic: a flexible trimmer for Illumina sequence data. Bioinformatics 30, 2114–2120.

Böndel KB, Lainer H, Nosenko T, Mboup M, Tellier A, Stephan W. 2015. North–South Colonization Associated with Local Adaptation of the Wild Tomato Species *Solanum chilense*. Molecular Biology and Evolution 32, 2932–2943.

Chaudhary A, Chen X, Gao J, Leśniewska B, Hammerl R, Dawid C, Schneitz K. 2020. The *Arabidopsisreceptor* kinase STRUBBELIG regulates the response to cellulose deficiency. PLOS Genetics 16, e1008433.

Coleman AD, Maroschek J, Raasch L, Takken FLW, Ranf S, Hückelhoven R. 2021. The *Arabidopsis*leucine-rich repeat receptor-like kinase MIK2 is a crucial component of early immune responses to a fungal-derived elicitor. New Phytologist 229, 3453–3466.

Cui J, Jiang N, Meng J, Yang G, Liu W, Zhou X, Ma N, Hou X, Luan, Y. 2016. LncRNA33732-respiratory burst oxidase module associated with WRKY1 in tomato-*Phytophthora infestansinteractions*. Plant Journal 97, 933–946.

Di X, Gomila J, Takken FLW. 2017. Involvement of salicylic acid, ethylene and jasmonic acid signalling pathways in the susceptibility of tomato to *Fusarium oxysporum*. Molecular Plant Pathology 18, 1024–1035.

Díaz J, Have A ten, Kan JAL van. 2002. The role of ethylene and wound signaling in resistance of tomato to *Botrytis cinerea*. Plant Physiology 129, 1341–1351.

Fischer I, Steige KA, Stephan W, Mboup M. 2013. Sequence evolution and expression regulation of stress-responsive genes in natural populations of wild tomato. PLOS ONE 8, e78182.

Freitas VS, Miranda R de S, Costa JH, Oliveira DF de, Paula S de O, Miguel E de C, Freire RS, Prisco JT, Gomes-Filho E. 2018. Ethylene triggers salt tolerance in maize genotypes by modulating polyamine catabolism enzymes associated with H2O2production. Environmental and Experimental Botany 145, 75–86.

Gravino M, Savatin DV, Macone A, De Lorenzo G. 2015. Ethylene production in *Botrytis cinerea-* and oligogalacturonide-induced immunity requires calcium-dependent protein kinases. Plant Journal 84, 1073–1086.

Halim VA, Eschen-Lippold L, Altmann S, Birschwilks M, Scheel D, Rosahl S. 2007. Salicylic acid is important for basal defense of *Solanum tuberosum* against *Phytophthora infestans*. Molecular Plant-Microbe Interactions 20, 1346–1352.

Hammond-Kosack KE, Silverman P, Raskin I, and Jones JDG. 1996. Race-specific elicitors of *Cladosporium fulvum* induce changes in cell morphology and the synthesis of ethylene and salicylic acid in tomato plants carrying the corresponding *Cf* disease resistance gene. Plant Physiology 110, 1381–1394.

Heitz T, Bergey DR, Ryan CA. 1997. A gene encoding a chloroplast-targeted lipoxygenase in tomato leaves is transiently induced by wounding, systemin, and methyl jasmonate. Plant Physiology 114, 1085–1093.

Herrera-Vásquez A, Salinas P, Holuigue L. 2015. Salicylic acid and reactive oxygen species interplay in the transcriptional control of defense genes expression. Frontiers in Plant Science 6.

Huang J, Zhao X, Bürger M, Wang Y, Chory J. 2021. Two interacting ethylene response factors regulate heat stress response. The Plant Cell 33, 338–357.

Jeun YC, Sigrist J, Buchenauer H. 2000. Biochemical and cytological studies on mechanisms of systemically induced resistance in tomato plants against *Phytophthora infestans*. Journal of Phytopathology 148, 129–140.

Kahlon PS, Seta SM, Zander G, Scheikl D, Hückelhoven R, Joosten MHAJ, Stam, R. 2020. Population studies of the wild tomato species *Solanum chilense* reveal geographically structured major gene-mediated pathogen resistance. Proceedings of the Royal Society B. 287, 20202723.

Kahlon PS, Stam R. 2021a. Polymorphisms in plants to restrict losses to pathogens: From gene family expansions to complex network evolution. Current Opinion in Plant Biology 62, 102040.

Kahlon PS, Stam R. 2021b. Protocol for chemiluminescence based detection of ROS production in tomato. protocols.io. DOI: 10.17504/protocols.io.beeejbbe

Kahlon PS, Verin M, Hückelhoven R, Stam R. 2021. Quantitative resistance differences between and within natural populations of *Solanum chilense* against the oomycete pathogen *Phytophthora infestans*. Ecology and Evolution 11, ece3.7610.

Kashyap SP, Kumari N, Mishra P, Prasad Moharana D, Aamir M, Singh B, Prasanna HC. 2020. Transcriptional regulation-mediating ROS homeostasis and physio-biochemical changes in wild tomato (*Solanum chilense*) and cultivated tomato (*Solanum lycopersicum*) under high salinity. Saudi Journal of Biological Sciences 27, 1999–2009.

Katz E, Li J-J, Jaegle B, Ashkenazy H, Abrahams SR, Bagaza C, Holden S, Pires CJ, Angelovici R, Kliebenstein DJ. 2021. Genetic variation, environment and demography intersect to shape *Arabidopsisdefense* metabolite variation across Europe. eLife 10, e67784.

Kim D, Langmead B, Salzberg SL. 2015. HISAT: a fast spliced aligner with low memory requirements. Nature Methods 12, 357–360.

Klarzynski O, Plesse B, Joubert J-M, Yvin J-C, Kopp M, Kloareg B, Fritig B. 2000. Linear β-1,3 Glucans are elicitors of defense responses in tobacco. Plant Physiology 124, 1027–1038.

Kruijt M, Kip DJ, Joosten MHAJ, Brandwagt BF, de Wit PJGM. 2005. The *Cf-4* and *Cf-9*resistance genes against *Cladosporium fulvum* are conserved in wild tomato species. Molecular Plant-Microbe Interactions 18, 1011–1021.

Liao Y, Smyth GK, Shi W. 2014. featureCounts: an efficient general purpose program for assigning sequence reads to genomic features. Bioinformatics 30, 923–930.

Liu Y, et al. 2020a. Diverse Roles of the Salicylic Acid Receptors NPR1 and NPR3/NPR4 in Plant Immunity. The Plant Cell 32, 4002–4016.

Liu H, Xue X, Yu Y, Xu M, Lu C, Meng X, Zhang B, Ding X, Chu Z. 2020b. Copper ions suppress abscisic acid biosynthesis to enhance defence against *Phytophthora infestans* in potato. Molecular Plant Pathology 21, 636–651.

Livak KJ, Schmittgen TD. 2001. Analysis of relative gene expression data using real-time quantitative PCR and the 2-ΔΔCT Method. Methods 25, 402–408.

Love MI, Huber W, Anders S. 2014. Moderated estimation of fold change and dispersion for RNA-seq data with DESeq2. Genome Biology 15, 550.

Marshall-Colón A, Kliebenstein, DJ. 2019. Plant Networks as Traits and Hypotheses: Moving Beyond Description. Trends in Plant Science 24, 840–852.

Meénard R, Alban S, de Ruffray P, Jamois F, Franz G, Fritig B, Yvin J-C, Kauffmann S. 2004. β-1,3 Glucan sulfate, but Not β-1,3 Glucan, induces the salicylic acid signaling pathway in Tobacco and *Arabidopsis*. The Plant Cell 16, 3020–3032.

Ng LM, Melcher K, Teh BT, Xu HE. 2014. Abscisic acid perception and signaling: structural mechanisms and applications. Acta Pharmacologica Sinica 35, 567–584.

Nie J, Yin Z, Li Z, Wu Y, Huang L. 2019 A small cysteine-rich protein from two kingdoms of microbes is recognized as a novel pathogen-associated molecular pattern. New Phytologist 222, 995–1011.

Nosenko T, Böndel KB, Kumpfmüller G, Stephan W. 2016. Adaptation to low temperatures in the wild tomato species *Solanum chilense*. Molecular Ecology 25, 2853–2869.

Ogran A, Landau N, Hanin N, Levy M, Gafni Y, Barazani O. 2016. Intraspecific variation in defense against a generalist lepidopteran herbivore in populations of *Eruca sativa*(Mill.). Ecology and Evolution 6, 363–374.

Peng J, Deng X, Jia S, Huang J, Miao X, Huang Y. 2004. Role of salicylic acid in tomato defense against cotton bollworm, *Helicoverpa armigera* Hubner. Zeitschrift für Naturforschung C 59, 856–862.

Pieterse CMJ, Leon-Reyes A, Van der Ent S, Van Wees SCM. 2009. Networking by small-molecule hormones in plant immunity. Nature Chemical Biology 5, 308–316.

R Core Team 2020. R: The R Project for Statistical Computing.

Raduski AR, Igić B. 2021. Biosystematic studies on the status of *Solanum chilense*. American Journal of Botany 108, 520–537.

Ramirez-Prado JS, Abulfaraj AA, Rayapuram N, Benhamed M, Hirt H. 2018. Plant immunity: from signaling to epigenetic control of defense. Trends in Plant Science 23, 833–844.

Riyazuddin R, Verma R, Singh K, Nisha N, Keisham M, Bhati KK, Kim ST, Gupta, R. 2020. Ethylene: a master regulator of salinity stress tolerance in plants. Biomolecules 10, 959.

Roberts R, Mainiero S, Powell AF, Liu AE, Shi K, Hind SR, Strickler SR, Collmer A, Martin GB. 2019. Natural variation for unusual host responses and flagellin-mediated immunity against *Pseudomonas syringae* in genetically diverse tomato accessions. New Phytologist 223, 447–461.

Sharaf EF, Farrag AA. 2004. Induced resistance in tomato plants by IAA against *Fusarium oxysporum lycopersici*. Polish Journal of Microbiology 53, 111–116.

Shibata Y, Kawakita K, Takemoto D. 2010. Age-related resistance of *Nicotiana benthamiana* against hemibiotrophic pathogen *Phytophthora infestans* requires both ethylene- and salicylic acid–mediated signaling pathways. Molecular Plant-Microbe Interactions 23, 1130–1142.

Smith JM, Heese A. 2014. Rapid bioassay to measure early reactive oxygen species production in *Arabidopsis* leave tissue in response to living *Pseudomonas syringae*. Plant Methods 10, 6.

Soltis NE, Atwell S, Shi G, Fordyce R, Gwinner R, Gao D, Shafi A, Kliebenstein DJ. 2019. Interactions of tomato and *Botrytis cinerea* genetic diversity: parsing the contributions of host differentiation, domestication, and pathogen variation. The Plant Cell 31, 502–519.

Song W, Ma X, Tan H, Zhou J. 2011. Abscisic acid enhances resistance to *Alternaria solani*in tomato seedlings. Plant Physiology and Biochemistry 49, 693–700.

Stam R, Scheikl D, Tellier A. 2017. The wild tomato species *Solanum chilense* shows variation in pathogen resistance between geographically distinct populations. PeerJ 5, e2910.

Stam R, Silva-Arias GA, Tellier A. 2019a. Subsets of *NLR* genes show differential signatures of adaptation during colonization of new habitats. New Phytologist 224, 367–379.

Stam R, Nosenko T, Hörger AC, Stephan W, Seidel M, Kuhn JMM, Haberer G, Tellier, A. 2019b. The *de Novo*reference genome and transcriptome assemblies of the wild tomato species *Solanum chilense* highlights birth and death of *NLR* genes between tomato species. G3: Genes, Genomes, Genetics 9, 3933–3941.

Steidele C. Stam R. 2021. Multi-omics approach highlights differences between functional RLP classes in *Arabidopsis thaliana*. BMC Genomics 22, 557.

Takemoto D, Shibata Y, Ojika M, Mizuno Y, Imano S, Ohtsu M, Sato I, Chiba S, Kawakita K, Rin S, Camagna M. 2018. Resistance to *Phytophthora infestans*: exploring genes required for disease resistance in Solanaceae plants. Journal of General Plant Pathology 84, 312–320.

Tian ZD, Liu J, Wang BL, Xie CH. 2006. Screening and expression analysis of *Phytophthora infestans* induced genes in potato leaves with horizontal resistance. Plant Cell Reports 25, 1094–1103.

Torres MA. 2010. ROS in biotic interactions. Physiologia Plantarum 138, 414–429.

Torres MA, Jones JDG, Dangl JL. 2006. Reactive oxygen species signaling in response to pathogens. Plant Physiology 141, 373–378.

Tziros GT, Samaras A, Karaoglanidis GS. 2021. Laminarin induces defense responses and efficiently controls olive leaf spot disease in olive. Molecules 26, 1043.

Ueeda M, Kubota M, Nishi K. 2005. Contribution of jasmonic acid to resistance against *Phytophthora*blight in *Capsicum annuum* cv. SCM334. Physiological and Molecular Plant Pathology 67, 149–154.

Van der Hoorn RAL, Kruijt M, Roth R, Brandwagt BF, Joosten MHAJ, Wit PJGMD. 2001. Intragenic recombination generated two distinct *Cf* genes that mediate AVR9 recognition in the natural population of *Lycopersicon pimpinellifolium*. Proceedings of the National Academy of Sciences of the United States of America 98, 10493–10498.

Van de Weyer A-L, Monteiro F, Furzer OJ, Nishimura MT, Cevik V, Witek K, Jones JDG, Dangl JL, Weigel D, Bemm F. 2019. A species-wide inventory of *NLR* genes and alleles in *Arabidopsis thaliana*. Cell 178, 1260–1272.e14.

VanderPlank JEV. 1963. Plant Diseases: Epidemics and Control. Academic Press.

Velásquez AC, Oney M, Huot B, Xu S, He SY. 2017. Diverse mechanisms of resistance to *Pseudomonas syringae* in a thousand natural accessions of *Arabidopsis thaliana*. New Phytologist 214, 1673–1687.

Von Kruedener S, Schempp H, Elstner EF. 1995. Gas chromatographic differentiation between myeloperoxidase activity and fenton-type oxidants. Free Radical Biology and Medicine 19, 141–146.

Wang L, et al. 2019. Arabidopsis *UBC 13* differentially regulates two programmed cell death pathways in responses to pathogen and low-temperature stress. New Phytologist 221, 919–934.

Wanke A, Rovenich H, Schwanke F, Velte S, Becker S, Hehemann J-H, Wawra S, Zuccaro A. 2020. Plant species-specific recognition of long and short β-1,3-linked glucans is mediated by different receptor systems. The Plant Journal 102, 1142–1156.

Windram O, Denby KJ. 2015. Modelling signaling networks underlying plant defence. Current Opinion in Plant Biology 27, 165–171.

Winkelmüller TM, et al. 2021. Gene expression evolution in pattern-triggered immunity within *Arabidopsis thaliana* and across Brassicaceae species. The Plant Cell, koab073.

Witek K, et al. 2021. A complex resistance locus in *Solanum americanum*recognizes a conserved *Phytophthora* effector. Nature Plants 7, 198–208.

Xin Z, Cai X, Chen S, Luo Z, Bian L, Li Z, Ge L, Chen Z. 2019. A disease resistance elicitor laminarin enhances tea defense against a piercing herbivore *Empoasca (Matsumurasca)*onukii Matsuda. Scientific Reports 9, 814.

Yang L, Zu Y-G, Tang Z-H. 2013. Ethylene improves *Arabidopsis* salt tolerance mainly via retaining K^+^ in shoots and roots rather than decreasing tissue Na^+^ content. Environmental and Experimental Botany 86, 60–69.

Yang X, Chen L, Yang Y, Guo X, Chen G, Xiong X, Dong D, Li G. 2020. Transcriptome analysis reveals that exogenous ethylene activates immune and defense responses in a high late blight resistant potato genotype. Scientific Reports 10, 21294.

Zhao Z et al. 2021. RPW8.1 enhances the ethylene-signaling pathway to feedback-attenuate its mediated cell death and disease resistance in *Arabidopsis*. New Phytologist 229, 516–531.

Zhu J-K. 2002. Salt and drought stress signal transduction in plants. Annual Review of Plant Biology. 53, 247–273.

