## Supplementary Figures for "Laminarin-triggered defence responses are geographically dependent for natural populations of *Solanum chilense*"

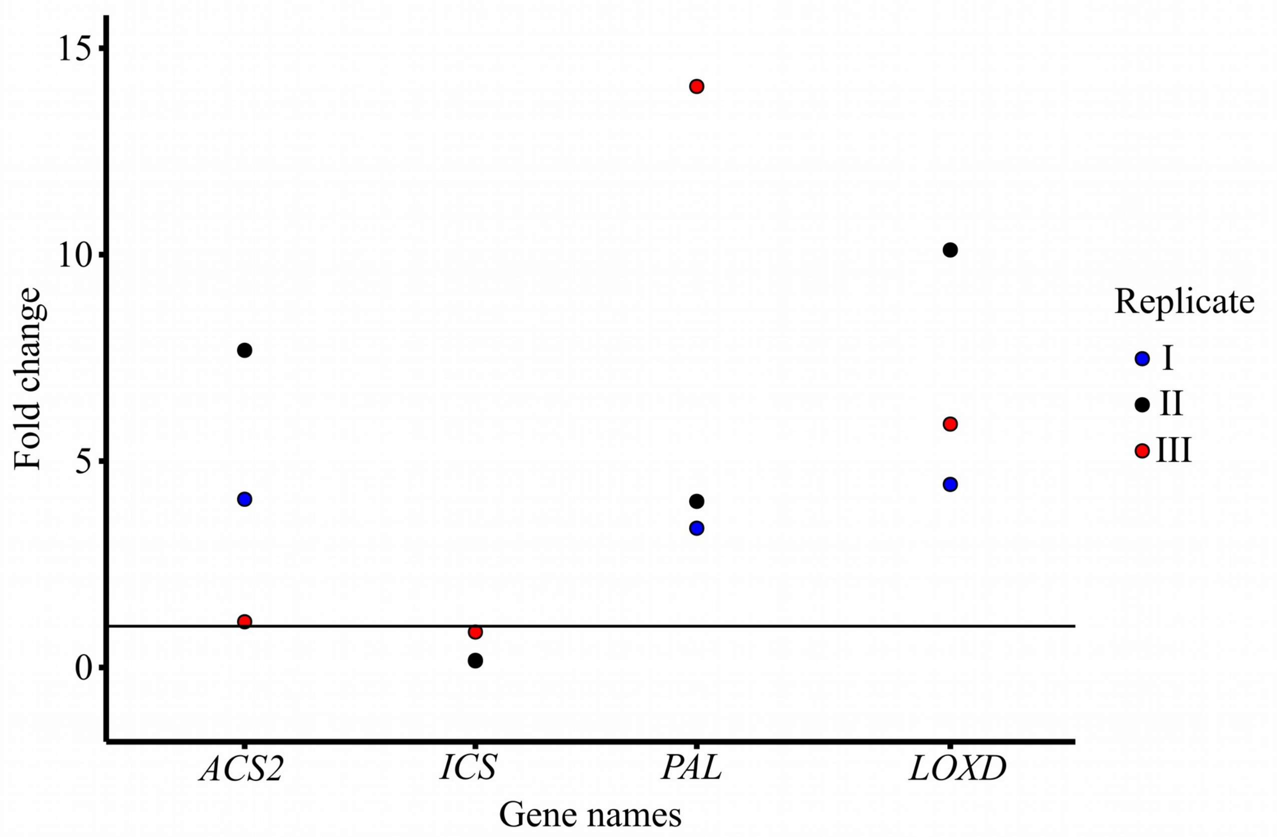


Figure S1: Expression level of key regulators of the ethylene (ET) production pathway, (*ACS2*), salicylic acid (SA) production pathway, (*ICS* & *PAL*) and jasmonic acid (JA) pathway, (*LOXD*) in *Solanum chilense* population LA1963 plant 02 upon 1.5 hours treatment with laminarin (1mg/ ml). Expression is shown in fold change (relative expression to non-treated samples), replicate indicates treatment performed on three different dates and each replicate represents average of three technical replicates calculated according to Livak and Schmittgen, (2001).

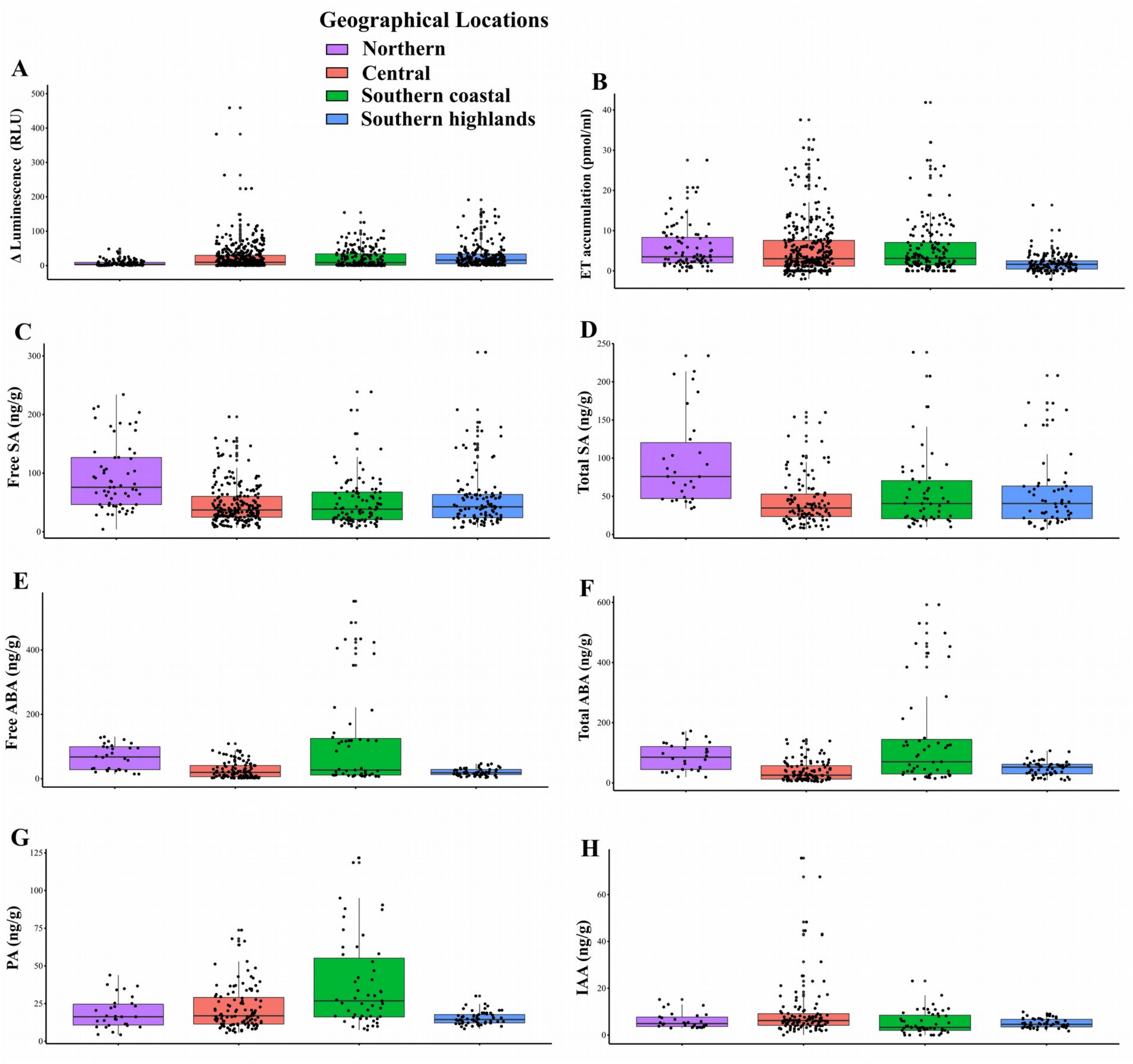


Figure S2: Geographical grouping for the measured components: level of ROS production and ET accumulation upon laminarin elicitation (a-b), basal levels of free SA (c), total SA (d), free abscisic acid ABA(e), total ABA(f), phaseic acid PA(g), and indoleacetic acid IAA (h). Each dot represents an individual measurement. The box plots summarize of three measurements per plant for each component from the respective geographical region. Color of box plot represents the geographical location.

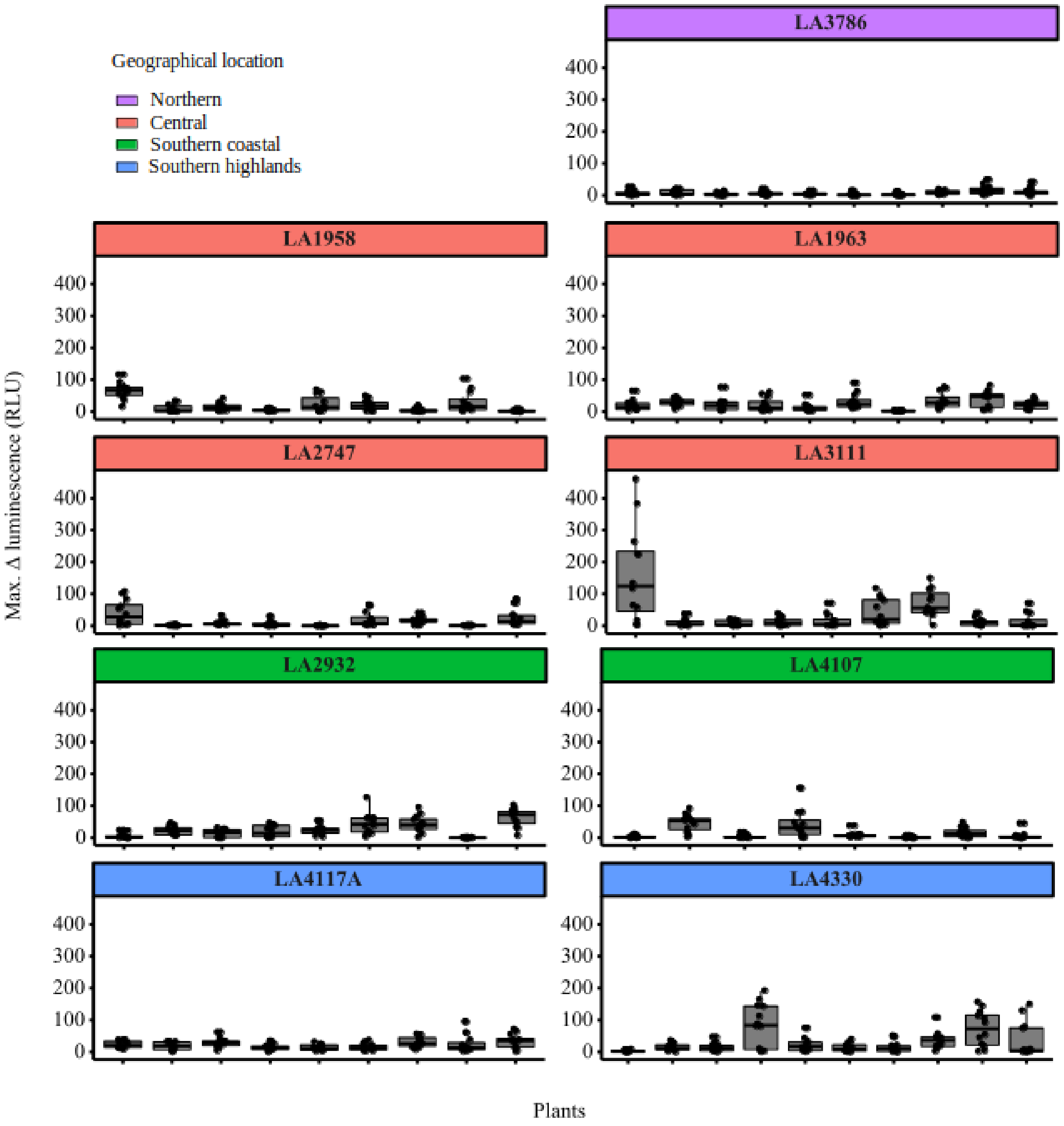


Figure S3 Maximum ROS accumulation in the leaf discs from *Solanum chilense* measured from 0-180 minutes upon elicitation with laminarin (1mg/ml). Each box plot represents an individual plant per population with data from one leaf disc represented as one data point accounting up to ten to twelve leaf discs per plant. Individual measurements were performed on three different dates (n=3-4 each date; 3x3(4)=10(12) leaf discs per plant). Y-axis shows relative luminescence unit (RLU). Each panel shows different population and colors represent the geographical location of the population.

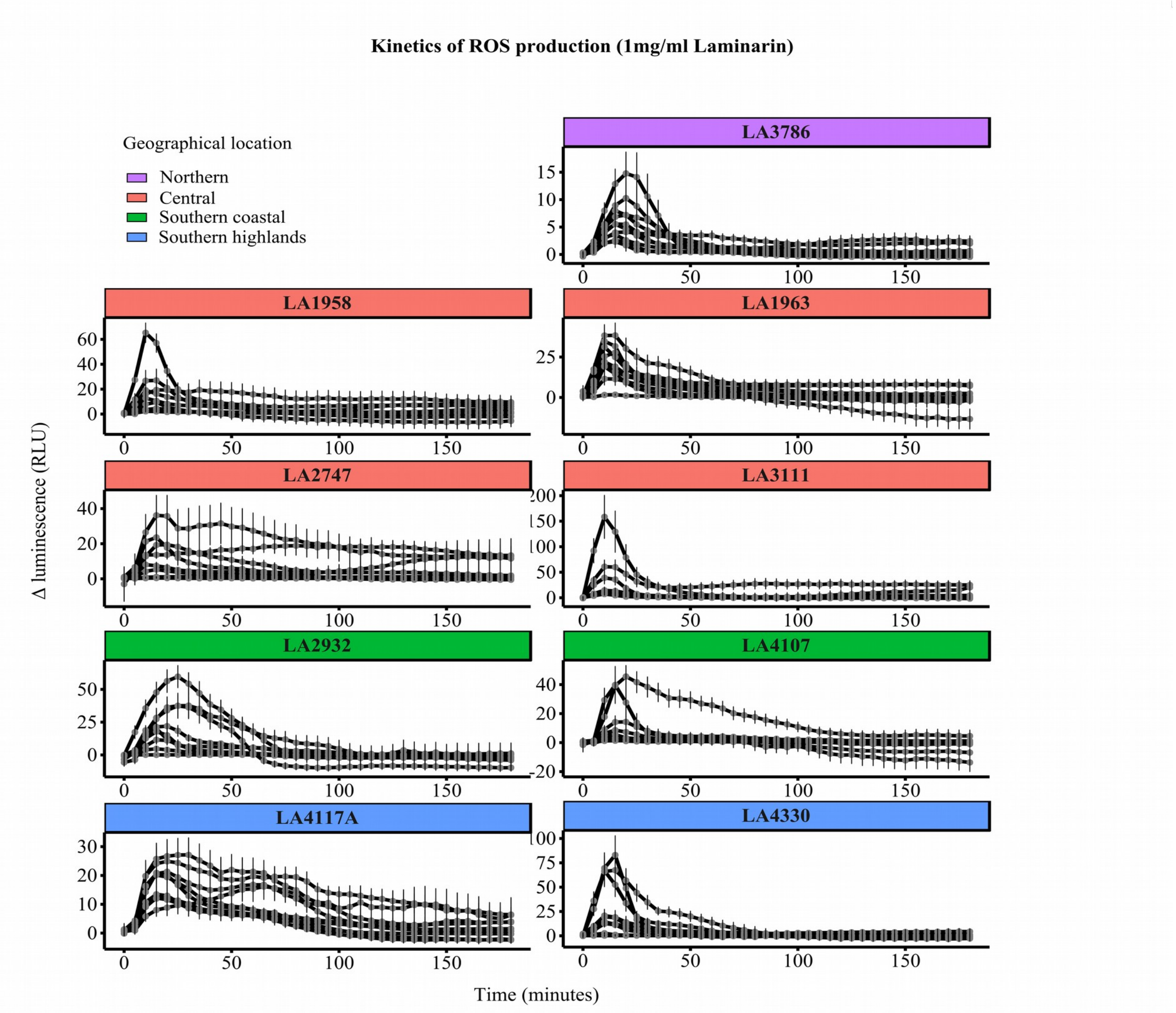


Figure S4: Kinetics of reactive oxygen species (ROS) production of *S. chilense* plants from 9 different populations (each facet represents one population and the color of the facet represents the geographical location of that population). Leaf discs were treated with 1mg/ml laminarin. Each line is the average mean of a single plant measured on three different dates with three to four technical replicates each (n =10(12), mean ± SE).

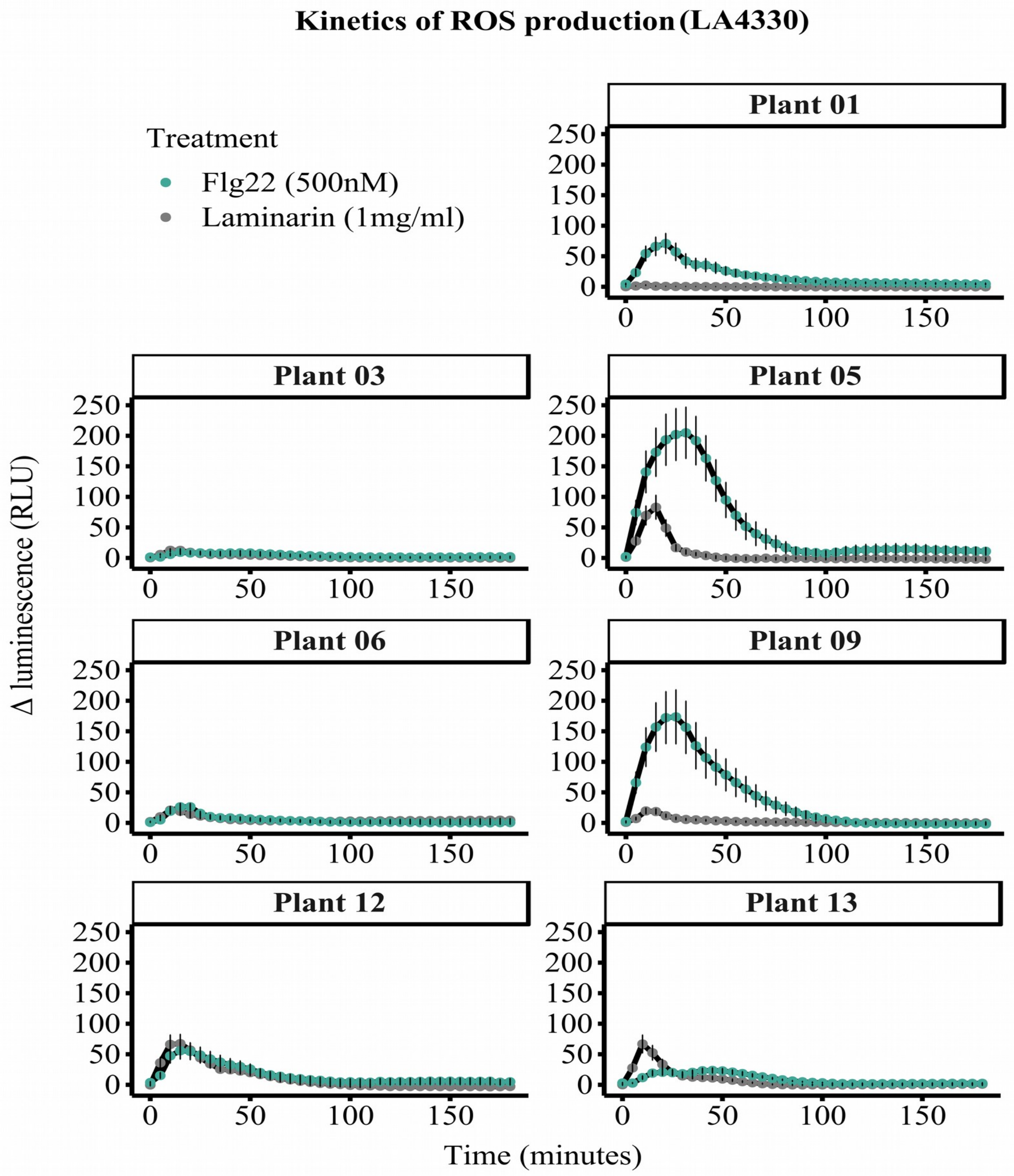


Figure S5: Kinetics of ROS production of *S. chilense* population LA4330 (7 plants) (each facet representis one plant) leaf discs treated with 1mg/ml laminarin (grey) or flg22 500nM (green). Each line is the average mean of a single plant measured on three different dates with three to four technical replicate each (n =10(12), mean ± SE).

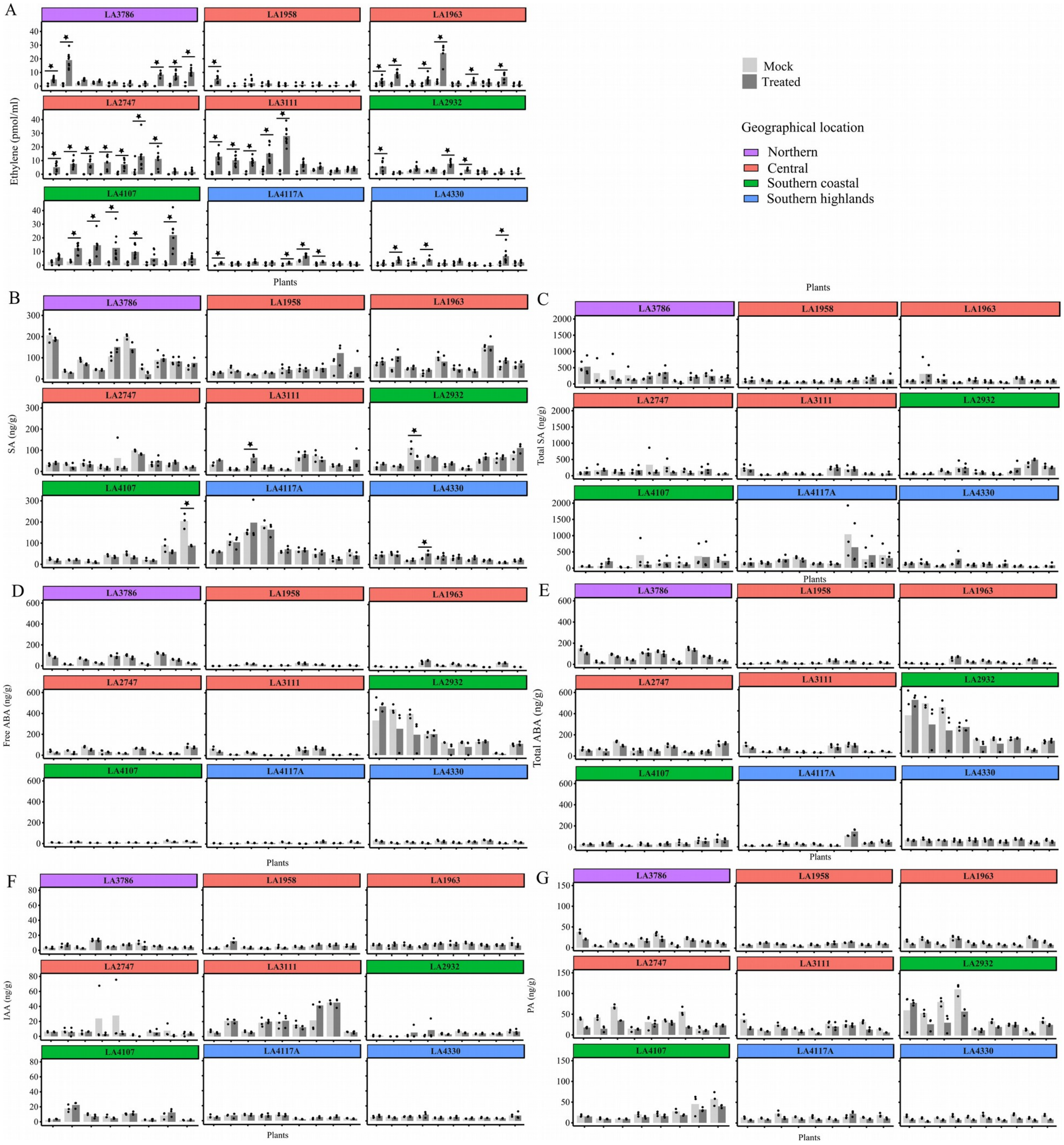


Figure S6: A) ET, B) free SA, C) total SA, D) free ABA, E) total ABA, F) IAA and G)PA levels (ng/g) in the leaf discs from 83 plants from *S. chilense* 3 hours upon elicitation with laminarin (1mg/ml), shown in dark grey and mock treatment (milliQ $H_20$), shown in light grey. Each bar pair represents a single individual from the population. Each bar is the mean of three data points which comprise of three sample measurements, each samples contained 50-100mg of fresh leaf weight obtained from the treated leaf discs (around 200 leaf discs per sample). Color of the facets represents the geographical location and each facet represents a single population.
